# Squamous-state excursions activate APOBEC3A in cancer

**DOI:** 10.64898/2026.05.24.727500

**Authors:** Josefine Striepen, Alexandra Dananberg, Aušrinė Ruzgaitė, Alyssa Hurley, Eléonore Toufektchan, Ashley Nichols, Hannah Rosenberg, Cameron Cordero, Tony M. Mertz, Roshan Xavier Norman, Richard Koche, Steven A. Roberts, John Maciejowski

## Abstract

The cytidine deaminase APOBEC3A is a major endogenous mutagen in human cancer, yet is rarely captured in bulk tumor RNA or protein profiles despite the prominent mutational scars it leaves in cancer genomes. The origin of these episodic mutational bursts has remained unclear, with prevailing models emphasizing sustained inflammatory signaling. Here, we show that APOBEC3A is induced in a rare subpopulation of cancer cells engaging a transient squamous differentiation program, linking APOBEC3A mutagenesis to lineage-state plasticity rather than persistent inflammatory signaling. Across breast and lung cancer cell lines and patient tumors, keratinocyte differentiation markers, including the stress keratins KRT6A and KRT16, are the strongest correlates of endogenous APOBEC3A expression. These days-long squamous-state excursions explain how APOBEC3A can leave durable mutational scars while remaining largely invisible to bulk tumor RNA and protein profiling. APOBEC3A catalytic activity reinforces selected components of this program through uracil excision and JNK-AP-1 signaling. The squamous differentiation transcription factor ZNF750 promotes APOBEC3A induction during squamous-state engagement in breast and lung cancer models. In established human squamous tumors, however, ZNF750 loss-of-function is associated with elevated APOBEC3A expression and APOBEC mutagenesis, revealing lineage-context-dependent regulation. These findings identify a transient differentiation state as a mutagenic intermediate, coupling cell-state plasticity to cancer genome evolution.

## INTRODUCTION

The APOBEC3 family of cytidine deaminases is among the most prevalent endogenous mutagens in human cancer, generating the SBS2 and SBS13 mutational signatures across breast, bladder, cervical, head and neck, lung, and esophageal tumors^1–5^. Ongoing APOBEC3 activity drives tumor heterogeneity, chromosomal instability, and acquired resistance to endocrine and targeted therapies, and is associated with poor clinical outcomes across diverse cancer types^6–10^. Human cancer cell line experiments have identified APOBEC3A as the dominant source of ongoing mutagenesis across most APOBEC-high lineages, despite APOBEC3A mRNA and protein accumulating to levels far below those of APOBEC3B in bulk measurements^4,11–14^. The disproportion between APOBEC3A expression and its mutational output remains a central unresolved feature of APOBEC biology in cancer.

APOBEC3A was identified as an interferon-stimulated gene prominently expressed in monocytes and macrophages, where it contributes to innate restriction of retroviruses and DNA viruses^15–18^. Prior work has identified several signals sufficient to induce APOBEC3A in defined experimental contexts, including interferon signaling through RIG-I/MAVS/IRF3, NF-κB-dependent induction by DNA damage, and EGFR inhibitor treatment in lung adenocarcinoma^19–21^. More recent studies have linked APOBEC3A to squamous differentiation, identifying GRHL3 as a transcriptional activator of APOBEC3A in differentiating squamous epithelia and squamous carcinomas^22^ and showing that mouse Apobec3 overexpression can promote squamous transdifferentiation through IL-1α/AP-1 signaling in a bladder cancer model^23^. These studies establish squamous differentiation as an APOBEC3A-associated context, but how APOBEC3A is induced endogenously in the non-squamous epithelial cancers where it dominates mutagenesis remains unknown.

APOBEC3A activity in cancer cells is episodic. Longitudinal mutation tracking in clonal cancer cell lines and phylogenetic analyses of human tumors have shown that APOBEC mutagenesis occurs in discrete bursts, with individual subclones acquiring APOBEC-signature mutations over short windows while related subclones acquire few or none^11–13,24,25^. This pattern suggests that APOBEC3A activity may be concentrated in rare cells or transient windows rather than reflected by stable bulk expression. The identity, duration, and reversibility of such an APOBEC3A-permissive state have remained unknown.

Here, we identify the cellular context in which APOBEC3A fires and define how a transient cell-state transition couples APOBEC3A activity to cancer genome evolution. Using a HaloTag knock-in reporter to isolate rare APOBEC3A-expressing cancer cells, we show that APOBEC3A, but not APOBEC3B, is selectively induced within a transient squamous differentiation program conserved across breast and lung cancer lines and patient tumors. APOBEC3A catalytic activity reinforces a subset of this program through uracil excision by UNG2 and JNK-AP-1 signaling. The transcription factor ZNF750 supports APOBEC3A expression during squamous-state entry in cancer cells. In established squamous tumors, however, ZNF750 loss-of-function is associated with elevated APOBEC3A and increased APOBEC mutagenesis, reflecting the dual activating and repressive functions of ZNF750 along the squamous differentiation trajectory. These findings identify a transient squamous differentiation state as a mutagenic intermediate, coupling cell-state plasticity to cancer genome evolution.

## RESULTS

### APOBEC3A activates in rare subclonal bursts in cancer cells

To examine where *APOBEC3A* is expressed in cancer, we analyzed CCLE bulk expression data across APOBEC-high tumor lineages^26^. *APOBEC3A* mRNA was consistently low, with distributions concentrated near the limit of detection, whereas *APOBEC3B* was expressed at substantially higher levels and across a wider dynamic range. This pattern was reproducible across all seven lineages examined and in a pooled set of 433 cell lines (Figure 1A). Quantitative RT-PCR across 34 breast cancer cell lines confirmed that *APOBEC3B* mRNA exceeded *APOBEC3A* by approximately 217-fold on average (Figure S1A), and *APOBEC3B* exceeded *APOBEC3A* in 95.7% of breast lines in DepMap (Figure S1B). Immunoblotting with an antibody recognizing both APOBEC3A and APOBEC3B^11^ confirmed this pattern at the protein level (Figure S1C). ssDNA deaminase assays across parental and knockout derivatives confirmed that APOBEC3B accounted for the majority of bulk activity, with APOBEC3A contributing a small but detectable share (Figures S1D and S1E). APOBEC3A is therefore expressed at very low bulk levels in cancer cell lines yet accounts for the majority of APOBEC-driven mutagenesis, deepening the paradox between its scarcity and its dominant mutational output.

**Figure 1.**
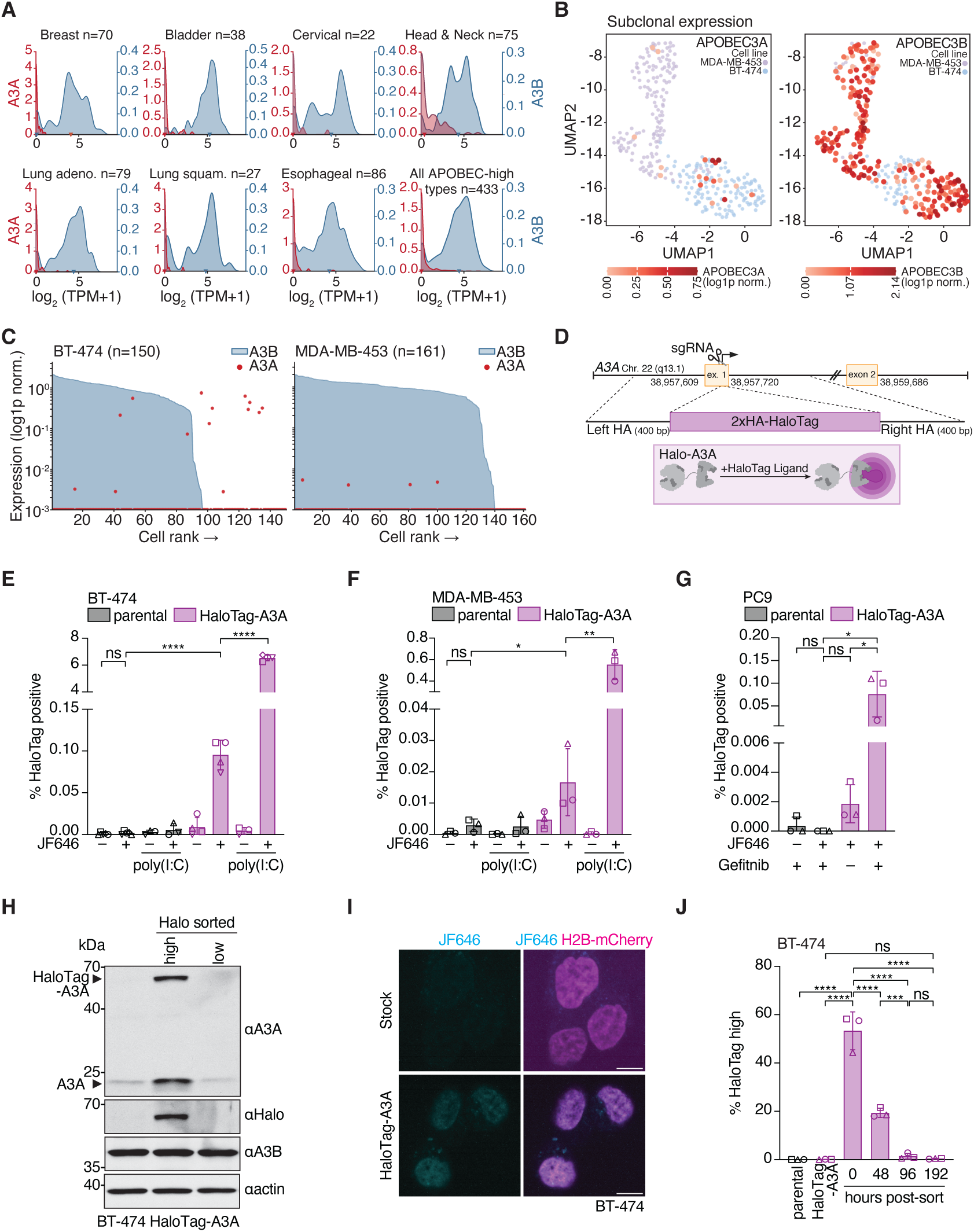
APOBEC3A is expressed in rare subclonal bursts in cancer cell lines and patient tumors. **(A)** Kernel density estimates of *APOBEC3A* (blue, left y-axis) and *APOBEC3B* (red, right y-axis) mRNA expression across APOBEC-high cancer cell line lineages (DepMap 24Q2). Expression as log₂(TPM+1). Y-axes show probability density, scaled independently for each paralog and lineage to visualize distribution shape. Arrowheads, median. *n* per lineage in panel titles. **(B)** Joint UMAP of BT-474 (*n*=314) and MDA-MB-453 (*n*=328) scRNA-seq, colored by *APOBEC3A* (left) or *APOBEC3B* (right) expression (log1p-normalized). **(C)** Per-cell expression rank plots for *APOBEC3A* (red dots) and *APOBEC3B* (blue shading) in BT-474 (*n* = 150) and MDA-MB-453 (*n* = 161) cells with detectable APOBEC3B expression. Red line, detection boundary (raw count ≥ 1). **(D)** Schematic of endogenous HaloTag-A3A knock-in at the *APOBEC3A* locus (chr22q13.1). **(E-G)** Percentage of HaloTag-A3A-positive cells. **(E)** BT-474, **(F)** MDA-MB-453 ±poly(I:C). **(G)** PC-9 ±gefitinib. *n*=3-4 replicates. Mean ± s.d. Two-tailed Student’s *t*-test; \*\*\**P* < 0.0001, \*\**P* < 0.01, \**P* < 0.05, ns = not significant. **(H)** Immunoblot of sorted HaloTag-high and HaloTag-low BT-474 HaloTag-A3A cells probed for APOBEC3A, HaloTag, APOBEC3B, and actin. Representative of *n* = 3 independent biological replicates. Antibodies listed in the methods. **(I)** Live-cell imaging of sorted HaloTag-A3A-positive BT-474 cells. JF646 (cyan), H2B-mCherry (magenta). Scale bars, 10 µm. **(J)** HaloTag-A3A-positive fraction at 0, 48, 96, and 192 h post-sort. Parental and unsorted HaloTag-A3A controls. *n* = 3 independent experiments. Mean ± s.d. One-way ANOVA with Tukey’s multiple comparisons. \*\*\*\**p* < 0.0001, \*\*\**p* < 0.001, ns = not significant. Individual replicate data points overlaid as distinct marker shapes (E, F, G, J).

To test whether heterogeneous *APOBEC3A* expression could explain the discrepancy between low bulk levels and a dominant role in mutagenesis, we performed single-cell RNA sequencing of BT-474 and MDA-MB-453. These lines depend on APOBEC3A for ongoing mutagenesis^11^ and rank in the top quartile for *APOBEC3A* expression among breast lines in DepMap (Figures S1A and S1B). *APOBEC3A* transcripts were detected in only a small fraction of cells in each line, while *APOBEC3B* was broadly expressed across the population (Figure 1B). Per-cell ranking of expression for each paralog confirmed this pattern. *APOBEC3B* was broadly expressed across the population, while *APOBEC3A* was detected above background in only a small fraction of cells (Figure 1C). Among other APOBEC3 paralogs detected in the dataset, *APOBEC3F* was broadly expressed in a manner similar to *APOBEC3B*, while *APOBEC3C*, *APOBEC3D*, and *APOBEC3G* showed sparse expression (Figure S1F).

In a published breast cancer scRNA-seq atlas (100,064 cells, 26 patients)^27^, APOBEC3A was detected in only 0.7% of cancer epithelial cells but in a substantially higher fraction of myeloid cells (Figures S1G and S1H), consistent with its established role as an interferon-stimulated gene in that lineage^16^. *APOBEC3A* expression in cancer cells is therefore restricted to a small subclonal population in both cell lines and patient tumors.

To confirm subclonal *APOBEC3A* expression at the protein level and to gain access to rare *APOBEC3A*-expressing cells for downstream characterization, we tagged endogenous APOBEC3A with a 2xHA-HaloTag cassette in BT-474 and MDA-MB-453 (Figure 1D, Figures S2A-C), enabling live-cell visualization and quantification of endogenous APOBEC3A protein by flow cytometry^28^. In BT-474 HaloTag-A3A cells, HaloTag-A3A was detected in approximately 0.1% of cells under baseline conditions and rose to approximately 6% following stimulation with the dsRNA mimetic poly(I:C), a known inducer of *APOBEC3A* expression (Figure 1E, Figure S2D)^19^. MDA-MB-453 HaloTag-A3A cells showed a similar pattern, with HaloTag-A3A detected in a small fraction of cells at baseline and increased upon poly(I:C) stimulation (Figure 1F, Figure S2E). PCR analysis of multiple subclones from a tagged MDA-

MB-453 line confirmed that the HaloTag cassette was present in all subclones examined, ruling out subclonal tag integration as a contributor to the rare HaloTag-A3A-positive populations (Figure S2F).

We extended this approach to PC-9, an EGFR-mutant lung adenocarcinoma line in which APOBEC3A is induced following EGFR inhibition^21^. Gefitinib induced APOBEC3A mRNA and protein without affecting APOBEC3B (Figures S2G and S2H), yet bulk deaminase activity was only minimally affected (Figure S2I), consistent with a small fraction of cells responding. Tagging endogenous *APOBEC3A* in PC-9 (Figure S2J) and treating with gefitinib induced a small HaloTag-A3A-positive population (0.02-0.1% across three replicates) (Figure 1G, Figure S2K), confirming the subclonal pattern in a third cancer lineage and in the context of drug-induced APOBEC3A activation.

Sorting HaloTag-A3A cells into high and low fractions confirmed that HaloTag-high BT-474 cells were enriched for both tagged and untagged APOBEC3A, while APOBEC3B was comparable between fractions (Figure 1H). The same pattern was observed in poly(I:C)-stimulated MDA-MB-453 HaloTag-A3A cells (Figure S2L) and in gefitinib-treated PC9 HaloTag-A3A cells (Figure S2M). In MDA-MB-453, IFIT1 was induced equally in both fractions (Figure S2L), indicating that all cells responded to poly(I:C) but only a small fraction translated this into APOBEC3A induction. RNA editing at the *DDOST* C558 site, an established cellular biomarker of APOBEC3A activity^29^, was elevated in HaloTag-high cells from both BT-474 and MDA-MB-453 (Figure S2N and S2O), further validating the HaloTag-A3A reporter as a marker for cells with active APOBEC3A. These results establish subclonal APOBEC3A expression at the protein level across multiple cancer lineages and validate the HaloTag-A3A reporter for prospective isolation of rare APOBEC3A-active cells.

Live-cell imaging of sorted HaloTag-A3A-positive BT-474 cells showed nuclear localization of HaloTag-A3A overlapping with the H2B-mCherry chromatin marker (Figure 1I). APOBEC3A nuclear localization was confirmed by subcellular fractionation in both BT-474 and MDA-MB-453 (Figures S3A and S3B). These observations resolve conflicting reports on APOBEC3A localization^30,31^ and place endogenous APOBEC3A in the compartment where it can access genomic DNA.

To test whether the A3A-high state is stable or transient, we sorted HaloTag-A3A-positive BT-474 cells and tracked them over 192 hours. Sorting enriched the HaloTag-positive fraction approximately 500-fold to 53%. The sorted population dropped to approximately 19% by 48 hours and approximately 1% by 96 hours, and was statistically indistinguishable from the unsorted starting population by 192 hours (Figure 1J). To exclude selective overgrowth of contaminating HaloTag-low cells, we confirmed that sorted HaloTag-high cells gave rise to colonies and expanded at rates comparable to HaloTag-low cells in both BT-474 and MDA-MB-453 (Figures S3C-H). Subclones derived from HaloTag-high cells expressed APOBEC3A across a range similar to HaloTag-low-derived subclones, rather than maintaining elevated expression (Figures S3I and S3J). These data establish that *APOBEC3A* expression bursts transiently in individual cells and resolves to baseline over a timescale of days, providing a cell-biological explanation for the episodic mutational pattern observed in cancer genomes^12^.

### APOBEC3A marks a squamous differentiation state in cancer cells

Interferon signaling has been proposed as a driver of *APOBEC3A* expression in cancer^15–17,19,20,32^, but quantitative RT-PCR of sorted BT-474 HaloTag-high cells showed no induction of *IFNB1* or *IFIT1*, despite robust *IFNB1* induction following poly(I:C) stimulation (Figure S4A). *ERBB2* mRNA was equivalent between fractions, excluding genetically distinct subclones (Figure S4A). To test the contribution of NF-κB signaling, an upstream activator of inflammatory *APOBEC3A* induction, we treated BT-474 cells with the IKK inhibitor TPCA-1. NF-κB inhibition had no effect on baseline APOBEC3A protein levels, while reducing poly(I:C)-induced IFIT1 expression in the same cells (Figure S4B). NF-κB inhibition similarly had no effect on the frequency of HaloTag-A3A high BT-474 cells (Figure S4C), indicating that the rare *APOBEC3A*-expressing population is not maintained by NF-κB signaling. Having found no evidence that interferon or NF-κB signaling drives the HaloTag-high state, we took an unbiased approach and sorted BT-474 HaloTag-high and HaloTag-low cells for bulk RNA sequencing (Figure 2A). The sort selectively enriched *APOBEC3A* without broadly inducing other *APOBEC3* family members (Figures S4D and S4E).

**Figure 2.**
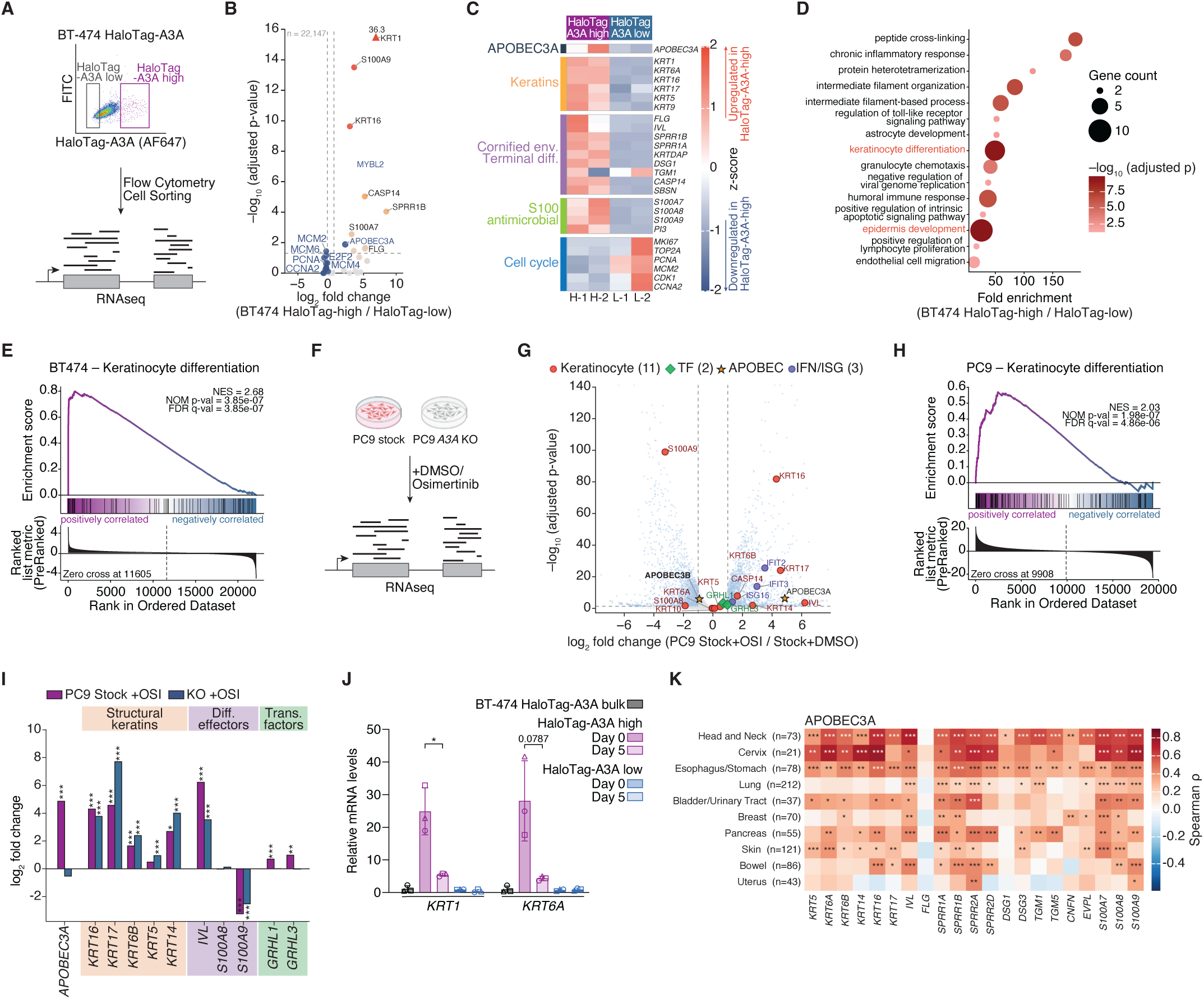
APOBEC3A expression marks a squamous differentiation state conserved across cancer lineages and patient tumors. **(A)** Schematic of the BT-474 HaloTag-A3A flow sort and bulk RNA sequencing workflow. **(B)** Volcano plot of BT-474 HaloTag-high vs HaloTag-low DEGs. Color coding in the panel. *n* = 2 per condition. DESeq2 Wald test, BH-FDR. Dashed lines, *p*adj = 0.05 and |log₂FC| = 0.5. **(C)** Heatmap of selected DEGs (row-wise z-score). Color scale clipped to [−2, +2]. **(D)** Gene ontology biological process enrichment of the 17 upregulated DEGs (*p*adj < 0.05, log₂FC > 1). Bubble size, gene count; color, −log₁₀(*p*adj). **(E)** GSEA of GO_KERATINOCYTE_DIFFERENTIATION across the ranked transcriptome. NES = 2.68, FDR *q* = 3.85 × 10⁻⁷. **(F)** Schematic of the PC-9 WT and *APOBEC3A* knockout osimertinib treatment and bulk RNA sequencing workflow. **(G)** Volcano plot of WT PC-9 osimertinib response. Color coding in the panel. DESeq2, BH-FDR. **(H)** GSEA of GO_KERATINOCYTE_DIFFERENTIATION in WT PC-9 osimertinib response. NES = 2.03, FDR *q* = 4.9 × 10⁻⁶. **(I)** Log₂FC of selected genes in WT (red) vs A3A-KO (blue) PC-9 osimertinib response. DESeq2. Significance stars, *p*adj from the corresponding contrast (\**p*adj < 0.05, \*\**p*adj < 0.01, \*\*\**p*adj < 0.001). **(J)** Quantitative RT-PCR of *KRT1* and *KRT6A* at day 0 and day 5 post-sort. Mean ± s.d. of *n* = 3 independent biological replicates. Statistical analysis by Welch’s two-sided two-sample *t*-test; \**p* < 0.05. Individual replicate data points overlaid as distinct marker shapes. **(K)** Heatmap of per-gene, per-lineage Spearman correlations between *APOBEC3A* expression and individual keratinocyte signature genes across DepMap cancer cell lines (24Q2). Lineages and *n* per lineage indicated. Significance: \**p* < 0.05, \*\**p* < 0.01, \*\*\**p* < 0.001.

Differential expression analysis identified a small but coherent set of transcripts that distinguished HaloTag-high from HaloTag-low cells (Figures 2B and 2C; Table S1). Of 22,147 genes tested, 17 were significantly upregulated and 17 downregulated (DESeq2, padj < 0.05). The upregulated set defined a squamous differentiation program spanning basal keratins (*KRT5*)^33,34^, stress keratins (*KRT6A*, *KRT16*, *KRT17*)^35–38^, the suprabasal keratin *KRT1*^39^, cornified envelope precursors (*SPRR1A*, *SPRR1B*, *IVL*, *FLG*)^40,41^, *TGM1*^42,43^, *CASP14*^44,45^, and *S100A7*/*A8*/*A9*^46^. This gene combination places HaloTag-high cells in an advanced squamous differentiation state resembling the keratinocyte activation program of wound healing and inflammatory skin disease^36,46–48^. *APOBEC3A* was upregulated in HaloTag-high cells (log^2^FC = 2.22, padj = 0.013), consistent with our prior immunoblotting and PCR data (Figure 1H). The downregulated set was enriched for cell cycle genes, including *MKI67*, *TOP2A*, *MCM2*, *PCNA*, *CCNA2*, and *CDK1*. The co-enrichment of markers from multiple epidermal layers^49–51^, together with downregulation of proliferative genes, is consistent with a disordered engagement of the squamous program characteristic of lineage plasticity in cancer^52–55^. Whether individual A3A-high cells simultaneously express markers from multiple epidermal layers or whether the sorted population captures cells at different positions along a squamous differentiation trajectory remains to be resolved at single-cell resolution. The transient transcriptional downregulation of cycling genes in freshly sorted HaloTag-high cells is reconcilable with normal long-term proliferative output (Figures S3C-E), because cells exit the A3A-high state within days and resume baseline cycling.

Gene ontology analysis of the 17 upregulated DEGs identified keratinocyte differentiation (padj = 2.77 × 10^-10^) and epidermis development (padj = 2.18 × 10^-10^) as the top enriched terms, confirmed by GSEA (NES = 2.68, FDR q = 3.85 × 10⁻⁷) (Figures 2D and 2E; Tables S2 and S3). ISG transcripts showed no differential expression in the RNA-seq data (Figure S4F), confirming the exclusion of interferon signaling. Quantitative RT-PCR validated upregulation of squamous markers in BT-474 HaloTag-high cells (Figure S4G), and a concordant pattern was observed in MDA-MB-453 HaloTag-high cells (Figure S4H).

To test whether the squamous program is a general correlate of endogenous *APOBEC3A* induction across cancer lineages and stimuli, we turned to PC-9 EGFR-mutant lung adenocarcinoma cells, in which *APOBEC3A* is induced during the drug-tolerant persister state following EGFR inhibition^21^. PC-9

WT and isogenic *APOBEC3A* knockout cells^13^ were treated with osimertinib or vehicle and profiled by bulk RNA sequencing (Figure 2F). Osimertinib engaged its canonical targets, suppressing EGFR-pathway feedback regulators and cell cycle drivers while upregulating alternative RTKs (Figure S4I). Against this backdrop, osimertinib induced a squamous differentiation program that strongly resembled the BT-474 HaloTag-high signature, including the basal and stress keratins *KRT5*, *KRT6A*, *KRT6B*, *KRT14*, *KRT16*, and *KRT17*, the cornified envelope precursors *IVL* and *CASP14*, and *APOBEC3A* itself (Figure 2G; Table S4). Gene set enrichment analysis confirmed strong enrichment for keratinocyte differentiation and epidermis development across the osimertinib-induced gene set (Figure 2H, Figure S4J; Table S3). *APOBEC3A* induction therefore coincides with a conserved squamous differentiation program across two cancer lineages and distinct induction stimuli.

To ask whether *APOBEC3A* is required for the osimertinib-induced squamous program, we compared the osimertinib response between WT and *APOBEC3A*-KO PC-9 cells. The bulk of the squamous program was induced to comparable levels in both genotypes, and GSEA confirmed strong enrichment for keratinocyte differentiation in the A3A-KO osimertinib response (Figure S4K). A subset of genes, however, showed asymmetric induction between genotypes. Most notably, the grainyhead-like transcription factors *GRHL1* and *GRHL3*, master regulators of epidermal differentiation^22,56^, were induced by osimertinib in WT but not A3A-KO cells (Figure 2I; Table S4). Quantitative RT-PCR across two independent A3A-KO clones and two WT subclones confirmed the loss of osimertinib-induced *GRHL3* expression in the absence of *APOBEC3A*, and a partial reduction in induction of *IVL* (Figure S4L). Together, these data indicate that APOBEC3A loss does not abolish the broader osimertinib-induced squamous program but selectively attenuates induction of a restricted subset of squamous differentiation genes, including GRHL3.

To test whether the squamous state reflects terminal differentiation or a reversible state, we sorted BT-474 HaloTag-high and HaloTag-low cells and measured squamous gene expression at day 0 and day 5 post-sort. Freshly sorted HaloTag-A3A-high cells expressed *KRT1* and *KRT6A* at 25- and 28-fold over unsorted parental cells, but returned to within approximately five-fold of baseline after five days in culture (Figure 2J). HaloTag-A3A-low cells showed no induction at either timepoint. The squamous state associated with endogenous *APOBEC3A* expression is therefore reversible and resolves on a timescale similar to HaloTag-A3A protein (Figure 1J), indicating a transient plastic state rather than commitment to terminal differentiation.

In TCGA data spanning 9,186 primary tumors, *APOBEC3A* expression correlated strongly with a 21-gene keratinocyte differentiation signature (Spearman ρ = 0.589, p < 10⁻¹⁶), significantly more so than *APOBEC3B* (ρ = 0.478, Meng-Rosenthal-Rubin p = 1.1 × 10⁻⁴⁰) (Figure S4M). At the gene level, *APOBEC3A* showed significant positive correlations with the majority of individual keratinocyte signature genes across every cancer type tested, while *APOBEC3B* did not (Figure S4N). Because bulk patient tumor expression can be confounded by infiltrating immune and stromal cells, we asked whether the same correlation held in cancer cell lines, where these confounds are absent. Across cancer cell lines from the DepMap database spanning 10 lineages, APOBEC3A expression correlated significantly with the keratinocyte signature at the per-gene level across squamous-enriched and non-squamous lineages (Figure 2K). *APOBEC3B* did not show this pattern in DepMap (Figure S4O; Meng-

Rosenthal-Rubin test for *APOBEC3A* vs *APOBEC3B* correlations, p < 10⁻¹⁵). To examine the association at single-cell resolution, we analyzed published patient scRNA-seq data from breast and lung adenocarcinoma^27,57^. *KRT16* was the top *APOBEC3A* correlate in both datasets, with keratinocyte and cornified envelope genes dominating the top-50 list (Figures S4P and S4Q). *APOBEC3B* correlates included cell cycle genes, with no overlap between the two paralog correlate lists. *GRHL1* ranked among the top 15 *APOBEC3A* correlates in the LUAD dataset (ρ = 0.118, FDR = 1.8 × 10⁻⁴⁰), independently validating the PC-9 knockout observations. Endogenous *APOBEC3A* expression therefore marks a squamous differentiation state conserved across cancer cell lines and patient tumors, a distinction not shared by *APOBEC3B*.

### APOBEC3A catalytic activity reinforces the squamous state

The A3A-dependent induction of a defined subset of the squamous program in PC-9 cells (Figure 2I) suggested that APOBEC3A may contribute to reinforcing the squamous state in which it is expressed. To test this directly, we introduced a doxycycline-inducible wild-type *APOBEC3A* construct into BT-474 cells and profiled the transcriptional response by bulk RNA sequencing. Dox induction elevated APOBEC3A substantially above endogenous Halo-high levels, providing a gain-of-function test. Dox induction produced a focused transcriptional response dominated by *APOBEC3A* itself (log^2^FC = 5.84, padj = 2 × 10⁻¹⁴⁵), accompanied by *KRT5*, *KRT6A*, *KRT1*, *CASP14*, *S100A8*, and *S100A9* (Figure 3A; Table S5). Gene ontology analysis confirmed keratinization as a top enriched term (Figures S5A and S5B).

**Figure 3.**
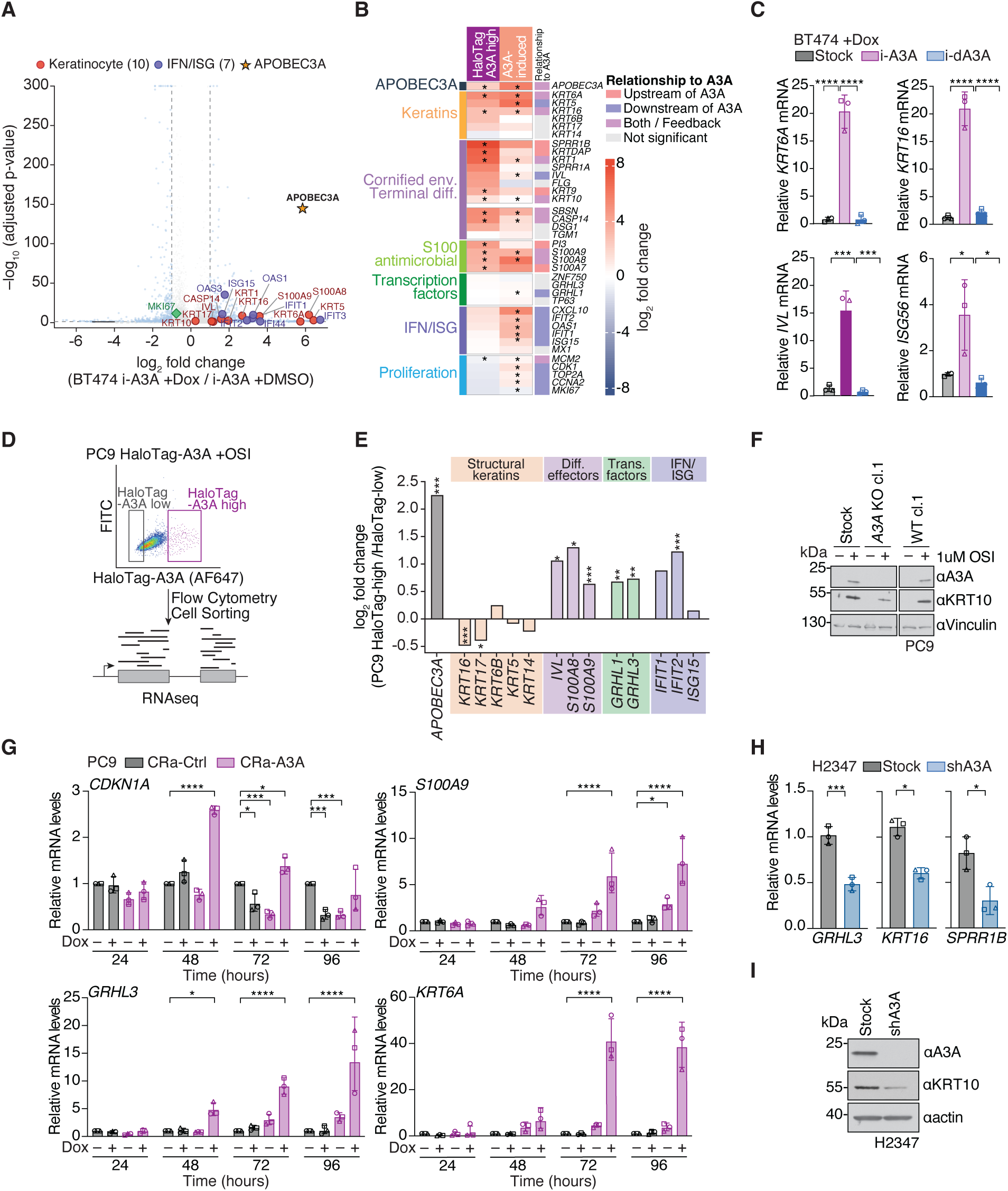
APOBEC3A catalytic activity engages a context-dependent subset of the squamous differentiation program. **(A)** Volcano plot of dox-inducible A3A vs uninduced BT-474 cells. *n* = 2 per condition. DESeq2, BH-FDR. Color coding in the panel. **(B)** Heatmap of log₂FC for selected genes in Halo-sort and Dox-OE contexts. *n* = 2 per condition. DESeq2. Asterisks, *p*adj < 0.05. Direction annotation classifies genes by significance in each context. **(C)** qPCR of selected genes in iA3A vs i-dA3A (E72Q) BT-474 cells. *n* = 3. One-way ANOVA with Tukey’s multiple comparisons. Error bars, mean ± s.d. \**p* < 0.05, \*\*\**p* < 0.001, \*\*\*\**p* < 0.0001. **(D)** Schematic of the PC-9 Halo-A3A osimertinib treatment, Halo-A3A-high and Halo-A3A-low FACS sort, and bulk RNA sequencing workflow. **(E)** Log₂FC of selected genes in PC-9 Halo-high vs Halo-low. *n* = 2 per condition. DESeq2, BH-FDR. Hatched bars, baseMean < 5. **(F)** Immunoblot of KRT10 and APOBEC3A in PC-9 WT and A3A-KO cells ± osimertinib. Representative of *n* = 2 replicates. **(G)** qPCR time course (24–96 h) in PC-9 dCas9-VPR cells with Ctrl sg or *A3A* sg ± doxycycline. *n* = 3. Two-way ANOVA with Dunnett multiple comparisons. Error bars, mean ± s.d. \**p* < 0.05, \*\*\**p* < 0.001, \*\*\*\**p* < 0.0001. **(H)** qPCR of *GRHL3*, *KRT16*, and *SPRR1B* in H2347 Stock vs shA3A. *n* = 3 biological replicates. Unpaired two-tailed *t*-test. Error bars, mean ± s.d. \**p* < 0.05, \*\*\**p* < 0.001. **(I)** Immunoblot of APOBEC3A, KRT10, and actin in NCI-H2347 Stock and shA3A cells. Representative of *n* = 1 biological experiment. Antibodies listed in the methods. Individual replicate data points overlaid as distinct marker shapes (C, G, and H).

Direct comparison of the Halo-sort and Dox-OE signatures revealed that A3A overexpression recapitulates only a subset of the endogenous state (Figure 3B, Figure S5C; Tables S1 and S5). Stress keratins (*KRT6A*, *KRT16*) and alarmins (*S100A8*, *S100A9*) were induced in both systems, marking the A3A-responsive arm. Terminal differentiation genes (*KRT1*, *SPRR1B*, *KRTDAP*, *KRT9*, *KRT10*) were strongly induced only in Halo-high cells. The proliferative suppression of Halo-high cells (*MKI67*, *CDK1*, *TOP2A*) was largely absent from Dox-OE. An ISG signature (*IFIT1*, *IFIT2*, *IFIT3*, *CXCL10*) was induced only in Dox-OE, consistent with supraphysiological A3A engaging pattern recognition pathways^58–60^. In contrast to wild-type iA3A, a dox-inducible catalytic-dead E72Q mutant (i-dA3A) failed to induce *KRT6A*, *KRT16*, *IVL*, or *ISG56* (Figure 3C), establishing that the transcriptional response requires A3A catalytic activity.

To test whether this extends to a second lineage, we sorted osimertinib-treated PC-9 Halo-A3A cells into high and low fractions for RNA sequencing (Figure 3D). APOBEC3A was enriched 2.2-fold in the Halo-high fraction (Table S4). The PC-9 pattern differed from BT-474 (Figure 3E; Figures S5D and S5E). Structural keratins (*KRT16*, *KRT17*) were not further enriched in Halo-high cells. Instead, *IVL*, *S100A8*, *S100A9*, and the squamous transcription factors *GRHL1* and *GRHL3* were significantly enriched, independently confirming A3A-associated *GRHL3* regulation (Figure 2I). Gene ontology analysis identified antimicrobial and innate defense terms (Figure S5F), but canonical ISGs were not enriched (Figure S5G), consistent with the absence of interferon signaling as a driver of endogenous A3A expression in this setting. KRT10 protein was attenuated in osimertinib-treated A3A-KO PC-9 cells relative to WT controls (Figure 3F), consistent with A3A reinforcing but not solely driving the squamous response.

CRISPR activation of endogenous APOBEC3A in PC-9 dCas9-VPR cells^61^ (Figure S5I-S5L) produced progressive upregulation of CDKN1A (48 h), followed by S100A9, GRHL3, and KRT6A (72-96 h) (Figure 3G). KRT6A reached approximately 40-fold induction, indicating that forced A3A expression engages structural keratins not coexpressed with endogenous A3A. The breadth of A3A-responsive genes therefore depends on cellular context and the level of A3A induction, with supraphysiological forcing engaging a wider program than endogenous levels drive.

In NCI-H2347 cells, which express the highest APOBEC3A among 212 lung cancer lines in DepMap (Figure S5H), shRNA-mediated A3A depletion reduced *GRHL3*, *KRT16*, and *SPRR1B* by approximately two-fold each (Figure 3H), confirming that endogenous A3A maintains squamous gene expression. A3A catalytic activity therefore engages a defined subset of the squamous program, and endogenous A3A is required to maintain this program in a naturally A3A-high cell line.

### APOBEC3A reinforces the squamous state through uracil excision and JNK-AP-1 signaling

The finding that A3A catalytic activity reinforces a specific subset of the squamous program (Figure 3) raised the question of how the cytidine deamination activity of A3A is translated into transcriptional consequences. A3A deaminates cytidine to uracil in single-stranded DNA, generating lesions that engage base excision repair and replication stress responses^62,63^. To identify the damage response kinases required for A3A-driven gene induction, we treated PC-9 dCas9-VPR cells with inhibitors of ATM or ATR during A3A induction and measured *SPRR1B* mRNA as a representative downstream readout (Figures S6A-D). *SPRR1B* induction was not blocked by ATM inhibition. ATR inhibition amplified *SPRR1B* approximately three-fold without affecting *A3A* mRNA levels, consistent with prior work showing that ATR restrains A3A-induced abasic site accumulation at replication forks and linking increased damage burden to increased transcriptional output^63,64^. These data exclude ATM and ATR as mediators of the A3A-driven response.

To ask whether base excision repair intermediates themselves transduce the A3A-driven transcriptional response, we examined BER pathway gene expression as a modifier of the A3A-keratinocyte association in TCGA patient tumors. Splitting 9,185 tumors by median expression of individual BER genes revealed that high expression of the nick-generating enzymes *UNG* and *PARP1* strengthened the A3A-keratinocyte correlation, while high expression of the nick-repair factors *XRCC1* and *LIG3* weakened it (Figure 4A). A composite BER imbalance score favoring nick generation over nick repair was associated with significantly stronger A3A-keratinocyte coupling (Δρ = +0.119, p = 1.75 × 10⁻¹⁸). These data predict that persistent BER intermediates transduce A3A catalytic activity into a transcriptional output.

**Figure 4.**
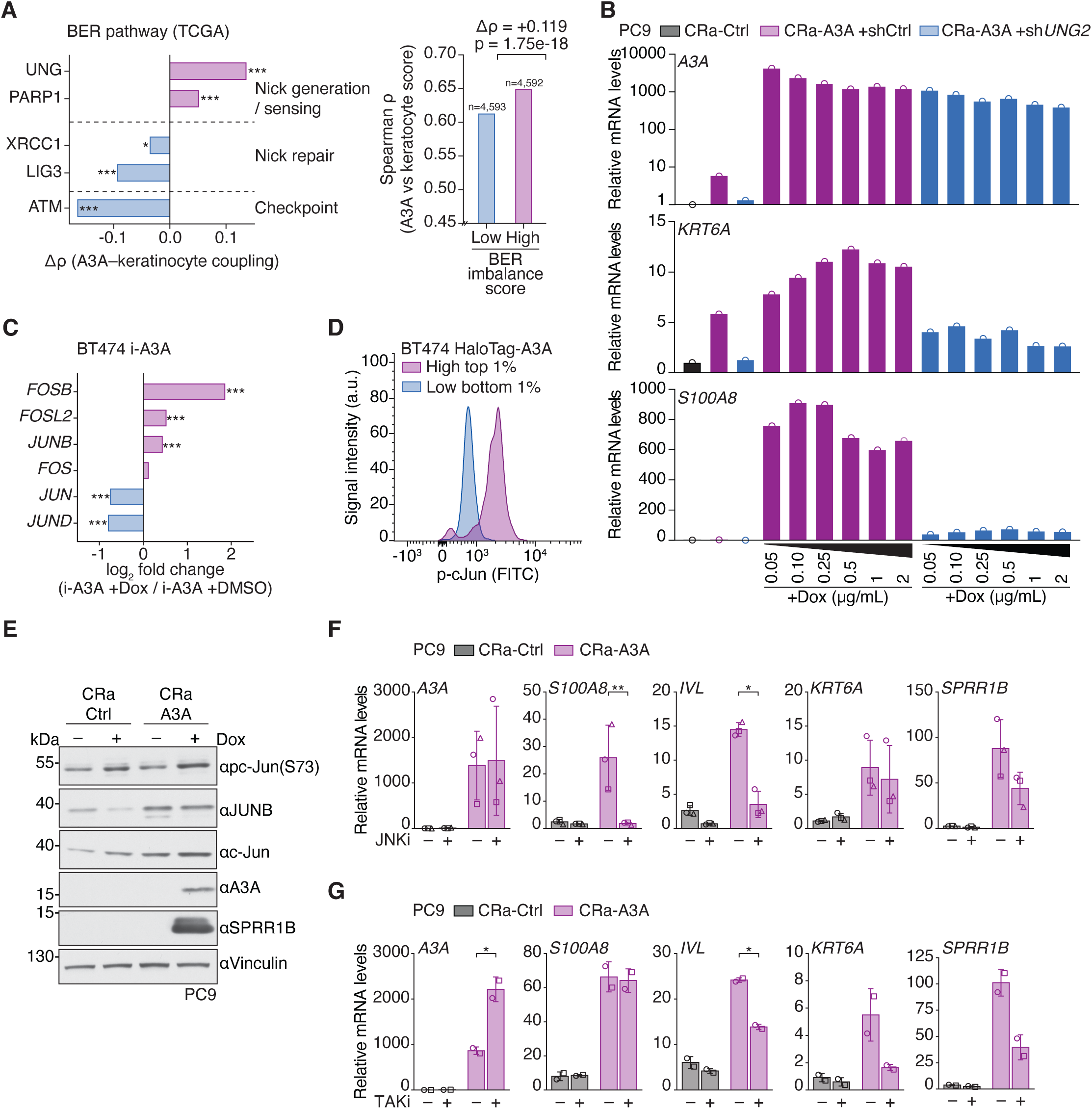
APOBEC3A reinforces the squamous state through uracil excision and JNK-AP-1 signaling. **(A)** BER pathway gene expression as a modifier of the *APOBEC3A*-keratinocyte differentiation association in TCGA (*n* = 9,185 primary tumors). Left, delta Spearman ρ (gene-high minus gene-low median split) for individual BER pathway genes, ordered by enzymatic step. Genes involved in nick generation and sensing (*UNG*, *PARP1*) strengthen the association. Genes involved in nick repair (*XRCC1*, *LIG3*) and checkpoint signaling (*ATM*) weaken it. Fisher z-test. Right, Spearman ρ between *APOBEC3A* and keratinocyte differentiation scores in tumors with low versus high BER imbalance composite scores (mean z-scored nick generation minus mean z-scored nick repair). \**p* < 0.05, \*\*\**p* < 0.001. **(B)** Quantitative RT-PCR of *APOBEC3A*, *KRT6A*, and *S100A8* in PC-9 dCas9-VPR cells expressing an *APOBEC3A*-targeting sgRNA with shCTRL or shUNG2 across a doxycycline titration (50–2000 ng/mL). *n* = 1, 72 hours post doxycycline treatment. **(C)** Bar plot of log₂ fold change for AP-1 transcription factor family members in BT-474 cells with doxycycline-inducible APOBEC3A relative to uninduced controls. *n* = 2 independent biological replicates per condition. DESeq2 Wald test with Benjamini-Hochberg adjustment. \*\*\**p*adj < 0.001. **(D)** Intracellular flow cytometry of phospho-c-Jun (Ser73) in BT-474 HaloTag-A3A-high (purple) and HaloTag-A3A-low (blue) cells. Representative of *n* = 2 independent experiments. **(E)** Immunoblot of phospho-c-Jun (Ser73), JUNB, c-Jun, APOBEC3A, SPRR1B, and vinculin in PC-9 dCas9-VPR cells expressing a control sgRNA (Ctrl sg) or an sgRNA targeting the *APOBEC3A* promoter (CRISPRa-A3A; CRa-A3A), with or without doxycycline. Representative of *n* = 2 independent biological replicates. Antibodies listed in the methods. **(F)** Quantitative RT-PCR of *APOBEC3A*, *S100A8*, *IVL*, *KRT6A*, and *SPRR1B* in PC-9 dCas9-VPR cells expressing Ctrl sg or *A3A* sg, treated with doxycycline (1 μg/mL) and/or JNK inhibitor (JNK-IN-8). *n* = 3 independent biological replicates. Welch’s two-sided *t*-test on log₂-transformed values comparing CRa-A3A DOX + JNKi vs CRa-A3A DOX. Error bars, mean ± s.d. \**p* < 0.05, \*\**p* < 0.01, \*\*\**p* < 0.001; ns, not significant. **(G)** Quantitative RT-PCR of *APOBEC3A*, *S100A8*, *IVL*, *KRT6A*, and *SPRR1B* in PC-9 dCas9-VPR cells expressing Ctrl sg or *A3A* sg, treated with doxycycline (1 μg/mL) and/or TAK1 inhibitor. *n* = 2 independent biological replicates. Welch’s two-sided *t*-test comparing CRa-A3A DOX + TAK1i vs CRa-A3A DOX. Error bars, mean ± s.d. \**p* < 0.05; ns, not significant. Individual replicate data points overlaid as distinct marker shapes (F and G).

To test this prediction directly, we depleted *UNG2* in PC-9 dCas9-VPR cells and induced A3A across a doxycycline titration series. At matched *A3A* expression levels, UNG2 depletion nearly abolished *S100A8* induction and attenuated *KRT6A* across all doses tested (Figure 4B), establishing uracil excision as a required step between A3A catalytic activity and the downstream transcriptional response.

With the requirement for uracil processing established, we asked which stress-responsive kinases act downstream to connect BER intermediates to the squamous transcriptional response. JNK is activated by DNA damage signaling through multiple pathways^65^ and the JNK-AP-1 axis is an established regulator of keratinocyte differentiation, with c-Jun directly regulating expression of cornified envelope precursors including SPRR family members and involucrin^66–68^. A3A induction in BT-474 reprogrammed AP-1 subunit expression, upregulating the differentiation-associated subunits FOSB, FOSL2, and JUNB and downregulating the proliferation-associated subunits JUN and JUND (Figure 4C; Table S5). To ask whether JNK-AP-1 activation occurs in cells with endogenous APOBEC3A expression, we measured phospho-c-Jun levels in BT-474 HaloTag-A3A cells by intracellular flow cytometry. HaloTag-A3A-high cells showed elevated phospho-c-Jun (Ser73) relative to HaloTag-A3A-low cells (Figure 4D), confirming that JNK-AP-1 signaling is activated in the endogenous APOBEC3A-high state. To validate this at the protein level in a second cancer lineage, we induced endogenous A3A in PC-9 cells using the dCas9-VPR CRISPR activation system and analyzed lysates by immunoblotting (Figure 4E). CRISPRa-mediated A3A induction was accompanied by phosphorylation of c-Jun on Ser73, a canonical JNK target site, with apparent increases in JUNB protein signal.

To ask whether JNK signaling contributes to A3A-driven gene induction, we treated PC-9 dCas9-VPR cells with the covalent JNK inhibitor JNK-IN-8^69^ during A3A induction and measured downstream gene expression by quantitative RT-PCR (Figure 4F). JNK inhibition did not affect A3A mRNA induction, placing JNK downstream of A3A transcription. S100A8 and IVL induction were significantly attenuated by JNK inhibitor (p < 0.01 and p < 0.05 respectively), while KRT6A induction was unaffected. SPRR1B showed a similar trend that did not reach statistical significance. To identify upstream activators contributing to JNK signaling in this context, we focused on TAK1, a MAP3K that activates JNK in response to cytokine and damage signaling and that is required for normal keratinocyte differentiation *in vivo*^70,71^. TAK1 inhibition partially attenuated A3A-induced IVL expression (p < 0.05), with similar trends for KRT6A and SPRR1B (Figure 4G). S100A8 induction was unaffected by TAK1 inhibition.

Together, these results establish a mechanistic relay in which A3A-generated uracils are excised by UNG and processed by base excision repair, generating persistent intermediates that activate JNK-AP-1 signaling and reinforce a subset of the squamous transcriptional program. The differential effects of UNG depletion, JNK inhibition, and TAK1 inhibition across the gene set indicate that multiple convergent and divergent signaling outputs downstream of uracil excision contribute to the broader A3A-driven transcriptional response.

### ZNF750 connects squamous differentiation to APOBEC3A induction

Having established that endogenous *APOBEC3A* marks a squamous differentiation state across cancer lineages, we sought to identify transcriptional regulators of the APOBEC3A-high state. We queried BT-474 Halo-high upregulated genes against 1,443 transcription factor perturbation signatures. The strongest enrichment was for genes downregulated after *ZNF750* knockdown in primary keratinocytes (adjusted p = 6.0 x 10^-15^; Figure 5A), with eleven of seventeen Halo-high DEGs overlapping ZNF750-activated differentiation targets. In osimertinib-treated PC-9 APOBEC3A-Halo-high cells, the strongest enrichments were *TP63* and *OVOL2* perturbation signatures (Figure 5A), placing endogenous *APOBEC3A* within a broader squamous transcription factor circuit across both lineages. By contrast, depletion of GRHL3, a reported activator of *APOBEC3A* in squamous epithelia^22^, did not reduce *APOBEC3A* expression in BT-474 cells despite efficient target knockdown (Figure S7A).

**Figure 5.**
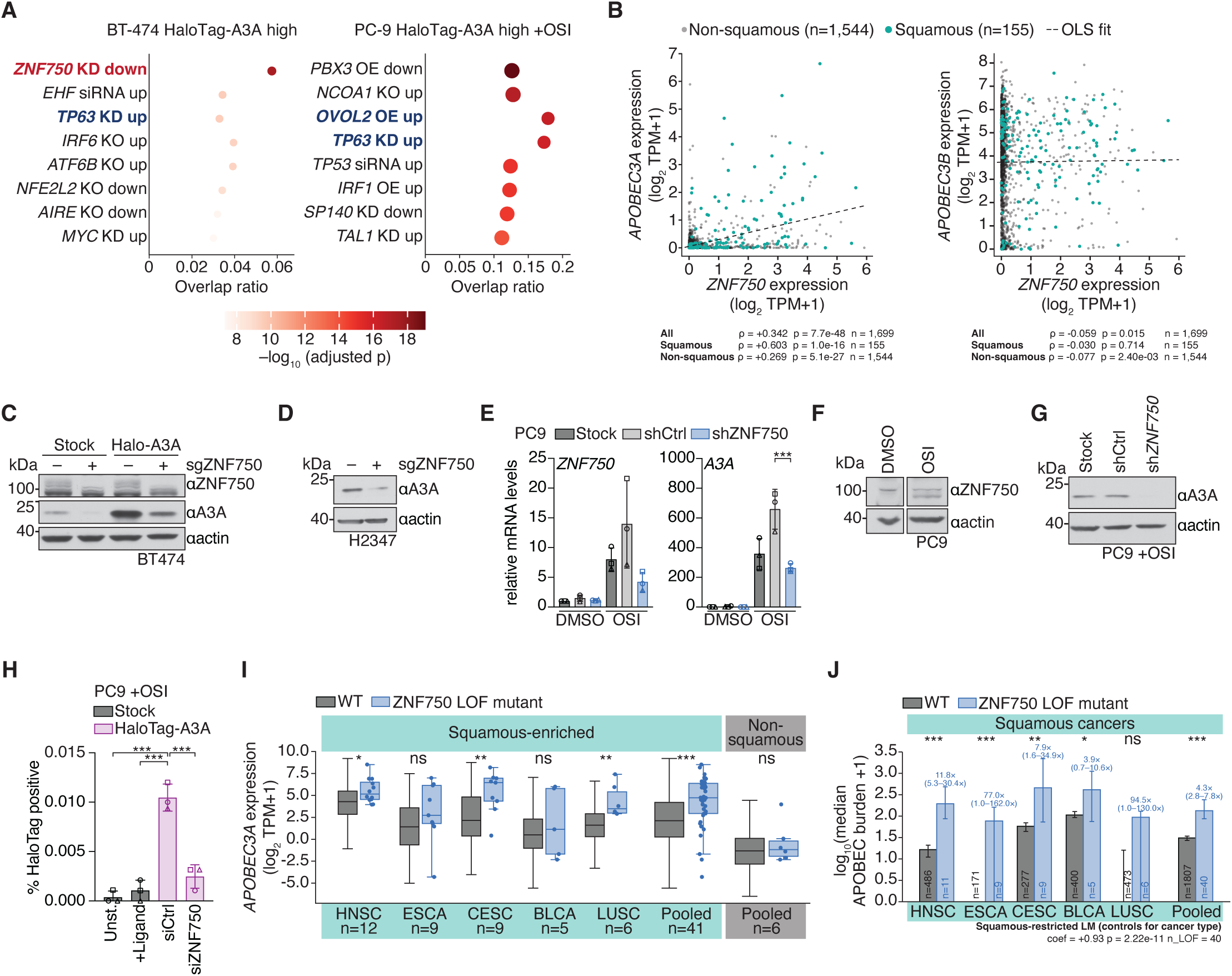
ZNF750 connects squamous differentiation to APOBEC3A mutagenesis. **(A)** Enrichr TF perturbation enrichment of BT-474 HaloTag-A3A-high DEGs (left) and osimertinib-treated PC-9 HaloTag-A3A-high DEGs (right). Top 8 hits by adjusted *p*-value. **(B)** Spearman correlation between *ZNF750* expression and *APOBEC3A* (left) or *APOBEC3B* (right) across 1,699 cancer cell lines (DepMap 24Q2). Expression as log₂(TPM+1). Squamous lineages (teal, *n* = 155) and non-squamous lineages (grey, *n* = 1,544) shown separately. Dashed line, OLS fit. Correlation statistics for all, squamous, and non-squamous subsets indicated. **(C–D)** Immunoblots following *ZNF750* depletion by sgRNA. **(C)** BT-474 stock and HaloTag-A3A cells probed for ZNF750, APOBEC3A, and actin. **(D)** NCI-H2347 cells probed for APOBEC3A and actin. Representative of *n* = 2 independent experiments. **(E)** Quantitative RT-PCR of *ZNF750* and *APOBEC3A* mRNA in PC-9 Stock, shCtrl, and shZNF750 cells treated with DMSO or 1 µM osimertinib. *n* = 3 independent biological replicates. Unpaired two-tailed *t*-test. Mean ± s.d. \**p* < 0.05, \*\**p* < 0.01, \*\*\**p* < 0.001. Individual replicate data points overlaid as distinct marker shapes. **(F)** Immunoblot of ZNF750 and actin in PC-9 cells treated with DMSO or 1 µM osimertinib. **(G)** Immunoblot of APOBEC3A and actin in osimertinib-treated PC-9 Stock, shCtrl, and shZNF750 cells. Representative of *n* = 2 independent biological replicates. Antibodies listed in the methods. **(H)** Percentage of HaloTag-A3A-positive cells in PC-9 HaloTag-A3A cells treated with 1 µM osimertinib following transfection with siCtrl or siZNF750. Stock and ligand-only controls shown. *n* = 3 independent technical replicates. Mean ± s.d. Individual replicate data points overlaid as distinct marker shapes. **(I)** *APOBEC3A* mRNA expression (log₂(TPM+1)) in *ZNF750* LOF versus WT tumors across five TCGA squamous-enriched cancer types and a pooled non-squamous cohort. Box plots show median, interquartile range, and 1.5× IQR whiskers. Individual LOF tumors shown as points. BH-adjusted Mann-Whitney U test. \**p* < 0.05, \*\**p* < 0.01, \*\*\**p* < 0.001, ns = not significant. *n* per group indicated. **(J)** APOBEC mutational burden (SBS2 + SBS13) in *ZNF750* LOF versus WT tumors across five TCGA squamous cancer types. Bars, median. Error bars, 95% bootstrap CI. Fold change and 95% CI indicated above each comparison. BH-adjusted Mann-Whitney U test. Pan-cancer pooled analysis by OLS linear regression controlling for cancer type (coefficient = +0.93, *p* = 2.22 × 10⁻¹¹, *n*_LOF = 40). \**p* < 0.05, \*\**p* < 0.01, \*\*\**p* < 0.001, ns = not significant.

ZNF750 is a zinc finger transcription factor that coordinates squamous terminal differentiation through interactions with KLF4 and chromatin regulators including KDM1A/LSD1-CoREST^72–74^. Across 1,699 cancer cell lines, *ZNF750* expression correlated positively with *APOBEC3A* but not *APOBEC3B* (Figure 5B). The top 100 *APOBEC3A*-correlated genes in patient tumor scRNA-seq were enriched for ZNF750-activated targets in both breast and lung adenocarcinoma datasets, while no other APOBEC3 paralog showed this enrichment (Figure S7B). To test whether ZNF750 supports APOBEC3A expression, we depleted *ZNF750* in breast and lung cancer cell lines. siRNA and sgRNA-mediated depletion reduced *APOBEC3A* mRNA and protein in BT-474 cells (Figure 5C; Figure S7C). sgRNA-mediated depletion similarly reduced APOBEC3A protein in NCI-H2347 cells (Figure 5D). In PC-9 cells, shZNF750 reduced *APOBEC3A* mRNA to approximately 15% of control levels (Figure S7D). Osimertinib induced *ZNF750* together with the squamous differentiation program at both the mRNA and protein levels (Figures 5E and 5F), and ZNF750 depletion attenuated osimertinib-induced APOBEC3A (Figures 5E and 5G). ZNF750 depletion also reduced the fraction of HaloTag-A3A-positive cells in osimertinib-treated PC-9 cells (Figure 5H), indicating that ZNF750 supports not only bulk APOBEC3A levels but also the frequency of cells entering the APOBEC3A-high state. Together, these data identify ZNF750 as a regulator of the transient APOBEC3A-high squamous state across breast and lung cancer models, connecting squamous differentiation transcription factor activity to the subclonal APOBEC3A bursts.

Integrative chromatin analysis further supported a regulatory relationship at the *APOBEC3A* locus. ZNF750 ChIP-seq peaks defined in differentiated keratinocytes overlapped promoter-proximal and distal *APOBEC3A*-associated elements that were accessible by ATAC-seq in BT-474, MDA-MB-453, and PC-9 cells, while published keratinocyte H3K27ac ChIP-seq showed reduced acetylation at these elements after ZNF750 depletion (Figure S7E). ZNF750 depletion had minimal effect on poly(I:C)-induced APOBEC3A or ISG56 mRNA or protein levels (Figure S7F and S7G), indicating that ZNF750 supports differentiation-associated but not acute poly(I:C)-driven induction of APOBEC3A.

This cancer-cell requirement contrasts with differentiated keratinocytes. Reanalysis of published ZNF750-depletion data from differentiated keratinocytes revealed that ZNF750 and KDM1A/LSD1 silence pattern recognition receptor genes including *TLR3*, *IFIH1*, and *DDX58*^72^. ZNF750 loss in this context derepresses these sensors, and poly(I:C) stimulation of ZNF750-depleted keratinocytes induced *APOBEC3A* 501-fold, the strongest response among APOBEC3 paralogs^72^. *APOBEC3A* was among genes both bound by ZNF750 and upregulated after ZNF750 loss in this system (Figure S7H). ZNF750 therefore has opposing effects on APOBEC3A depending on whether it operates in the context of squamous-state entry or in cells that have completed squamous differentiation.

ZNF750 is recurrently mutated in squamous cancers, and we therefore asked whether ZNF750 loss has a distinct relationship with APOBEC3A in squamous tumors, where squamous lineage identity is already established. Across TCGA squamous and squamous-lineage-enriched cancer cohorts, tumors carrying *ZNF750* loss-of-function mutations showed elevated *APOBEC3A* mRNA relative to wild-type tumors (pan-squamous pooled p = 4.5 x 10^-7^), with significant effects in HNSC, CESC, and LUSC (Figure 5I). *ZNF750* loss-of-function mutations were concentrated in squamous cancer types and rare outside this context. *APOBEC3B* was not consistently elevated (Figure S7I). Consistent with elevated *APOBEC3A* expression, SBS2 and SBS13 mutations were increased in *ZNF750* loss-of-function tumors across squamous cancers in a model controlling for cancer type and total non-APOBEC mutational burden, which was not elevated in *ZNF750* loss-of-function tumors (Figure S7J; model-estimated fold change = 8.4, 95% CI = 4.5 to 15.7, p = 2.22 × 10⁻¹¹, n = 40 LOF tumors among 1,847 squamous-lineage tumors; Figure S7J). The association was robust to leave-one-cancer-type-out analyses, including removal of ESCA, the only cohort with prior reports linking *ZNF750* mutations to APOBEC mutagenesis (Methods)^75–77^. Only 3 of 41 *ZNF750* loss-of-function events occurred in APOBEC trinucleotide contexts, and 52% were frameshift indels that are unlikely to be direct products of APOBEC cytidine deamination (Figure S7K), arguing against direct generation of most *ZNF750* loss-of-function mutations by APOBEC mutagenesis.

These data provide a lineage-context framework for the previously observed association between *ZNF750* mutations and APOBEC mutational signatures in esophageal squamous cell carcinoma^75–77^. These data place ZNF750 at an interface between squamous differentiation and APOBEC mutagenesis, with opposing net effects on APOBEC3A depending on whether cells are entering or have completed the squamous program. ZNF750 supports APOBEC3A induction during squamous-state entry in breast and lung cancer models, whereas *ZNF750* loss-of-function in established squamous tumors is associated with elevated *APOBEC3A* expression and increased SBS2/SBS13 mutagenesis (Figure S7L).

## DISCUSSION

Our findings recast APOBEC3A mutagenesis in cancer as a consequence of transient squamous differentiation rather than sustained inflammatory signaling. Across breast and lung cancer cell lines, APOBEC3A is selectively induced in cells engaging a transient squamous differentiation program, and keratinocyte differentiation markers are the strongest correlates of APOBEC3A in patient tumor single-cell data. These days-long squamous-state excursions resolve over days, returning the APOBEC3A-high fraction to its rare baseline frequency, and during the window of expression, APOBEC3A catalytic activity reinforces a subset of the squamous program in part through JNK/AP-1 signaling. ZNF750 supports the squamous-state circuitry in which APOBEC3A induction occurs. The result is a transient differentiation state that functions as a mutagenic intermediate, coupling cell-state plasticity to cancer genome evolution.

This excursion model provides a cell-state-level explanation for observed patterns of episodic APOBEC mutagenesis in cancer genomes^11,12,78^. Subcellular fractionation confirms that endogenous APOBEC3A concentrates in the nucleus during the squamous-state excursion, and *DDOST* C558 RNA editing is elevated in sorted APOBEC3A-high cells relative to APOBEC3A-low cells, confirming that higher APOBEC3A protein corresponds to higher deaminase activity. The squamous excursion positions active APOBEC3A in the compartment where genomic DNA is accessible. A cell that enters the squamous-like state is therefore positioned to acquire APOBEC3A-driven mutations during the window of activity and to exit the state within days, leaving durable mutational scars while rendering APOBEC3A largely invisible to bulk RNA and protein profiling. This association between APOBEC3A and the squamous program is not shared by APOBEC3B. APOBEC3B, despite higher bulk expression, does not show the same keratinocyte-dominated correlation pattern across DepMap or patient single-cell datasets, reinforcing the paralog-specific biology that underlies APOBEC3A’s dominant but cryptic mutagenic role.

APOBEC3A was originally identified as phorbolin-1, a transcript upregulated in psoriatic keratinocytes^79,80^. GRHL3 is a transcriptional activator of APOBEC3A during keratinocyte differentiation^22^, although GRHL3 depletion did not reduce APOBEC3A in BT-474 cells, indicating that GRHL3 is not universally required for APOBEC3A induction across cancer lineages. In a mouse urothelial carcinoma model, forced overexpression of Apobec3, an ortholog most similar to human APOBEC3F with considerably lower deaminase activity than APOBEC3A, drove squamous transdifferentiation through IL-1α and AP-1^23^. Our work demonstrates that endogenous human APOBEC3A engages a similar program at physiological expression levels in a transient, reversible cell state. The conservation of APOBEC3A within the squamous differentiation program raises the possibility that APOBEC3A plays a functional role during normal keratinocyte differentiation, potentially through the same uracil-dependent transcriptional mechanism described here. In normal tissue, terminal differentiation would provide a natural containment strategy, restricting a potent mutator to cells exiting the proliferative compartment.

UNG2 depletion abolished the A3A-driven transcriptional response, establishing uracil excision as an obligate step in a relay connecting A3A catalytic activity to transcriptional reinforcement of the squamous program. In patient tumors, BER pathway gene expression modified the strength of the A3A-keratinocyte association in a manner consistent with persistent repair intermediates, rather than completed repair, transducing the signal. ATR inhibition amplified rather than blocked the downstream response, consistent with increased unrepaired lesion burden driving greater transcriptional output^62–64^. TAK1 inhibition attenuated selected A3A-induced outputs, positioning TAK1 as a candidate kinase connecting BER intermediates to JNK-AP-1 activation. The molecular steps between uracil excision and JNK phosphorylation remain to be fully defined.

The effect of ZNF750 on APOBEC3A depends on position along the squamous differentiation trajectory. In cancer cells adopting a squamous-like program, ZNF750 depletion reduces *APOBEC3A* expression across breast and lung cancer models, and integrative chromatin analysis supports a regulatory relationship at the *APOBEC3A* locus. ZNF750 ChIP-seq peaks defined in differentiated keratinocytes overlap promoter-proximal and distal elements that are accessible in BT-474, MDA-MB-453, and PC-9 cells, and ZNF750 depletion reduces H3K27ac at these elements. A candidate ZNF750 binding motif is present at the APOBEC3A promoter within this element, consistent with direct transcriptional regulation. This positive relationship is consistent with the established role of ZNF750 as a transcriptional activator of differentiation genes in normal squamous epithelia^73,74^. In differentiated keratinocytes, ZNF750 also represses a distinct set of targets including pattern recognition receptor genes through KDM1A/ CoREST-mediated silencing, and *APOBEC3A* is among the genes derepressed after ZNF750 loss in that context^72^.

In established human squamous tumors, where squamous identity is already established, ZNF750 loss-of-function is associated with elevated *APOBEC3A* expression and increased SBS2/SBS13 mutational burden, consistent with loss of the repressive arm. These opposing relationships can be reconciled by a model in which ZNF750 operates at different points along the squamous differentiation trajectory. In non-squamous cancer cells, ZNF750 supports initial engagement of the squamous program and the APOBEC3A induction that accompanies it. In tissues that have completed squamous differentiation, ZNF750 restrains APOBEC3A as part of its broader silencing of innate immune effectors. These findings provide a lineage-based framework for the previously unexplained association between *ZNF750* mutations and APOBEC mutational signatures in esophageal squamous cell carcinoma^75–77^. More broadly, squamous reprogramming is a recurrent feature of lineage plasticity in metastatic colorectal cancer^81,82^, and the squamous excursion described here provides a candidate mechanism linking APOBEC3A activity to the plastic states reported in metastatic disease.

The subclonal nature of APOBEC3A expression has implications for therapeutic strategies that seek to exploit APOBEC activity. Synthetic lethality approaches have been developed using constitutive overexpression systems that model a state only a rare subpopulation occupies endogenously^62,63,83–86^. Therapeutic windows *in vivo* may therefore be narrower than these models predict, because synthetic lethal interactions would target only the rare A3A-expressing fraction rather than the tumor bulk. APOBEC mutations have also been proposed as a source of tumor neoantigens for immunotherapy^87^. In tumors with ongoing subclonal APOBEC activity, the bursting pattern described here predicts that APOBEC-generated neoepitopes will be distributed across many small subclones rather than shared across the tumor bulk. Because subclonal neoantigens elicit weaker immune responses than clonal neoantigens^88^, the immunotherapeutic potential of ongoing APOBEC mutagenesis may be more limited than models based on bulk mutational burden imply.

These findings identify a transient differentiation state as a mutagenic intermediate, coupling cell-state plasticity to cancer genome evolution. We show that a rare, reversible, prospectively isolatable cell state can explain why a nearly invisible transcript dominates mutational output across APOBEC-high epithelial cancers. The squamous excursion model reframes APOBEC3A in epithelial cancers not as an aberrantly activated immune effector but as a component of a lineage program that cancer cells transiently engage, with durable consequences for the mutational landscape.

### Limitations of the study

Several aspects of the model presented here require additional investigation. First, the molecular relay connecting APOBEC3A catalytic activity to JNK-AP-1 signaling is incompletely defined. We establish UNG2 as required for the transcriptional response, and JNK and TAK1 inhibition attenuates a subset of A3A-induced squamous genes, but the specific BER intermediates and signaling steps linking uracil excision to JNK activation remain to be identified. Second, the upstream signals that initiate squamous-state engagement in cancer cells are unknown. The transience and rarity of the state suggest a cell-intrinsic trigger that we have not yet defined. Third, while ZNF750 ChIP-seq peaks from differentiated keratinocytes overlap accessible chromatin at the APOBEC3A locus in our cancer cell lines and a candidate ZNF750 binding motif is present in this element, direct ZNF750 binding at the APOBEC3A locus in cancer cells has not been confirmed by ChIP. The regulatory relationship is therefore consistent with direct transcriptional control but does not formally establish it. Finally, our cell-biological characterization of the squamous excursion is performed in breast and lung cancer models. Extension to additional APOBEC-high cancer lineages including bladder, head and neck, cervical, and esophageal cancers will require further work, though our patient genomic analyses suggest the model is likely to generalize across these contexts.

## Reagents and Resources Table

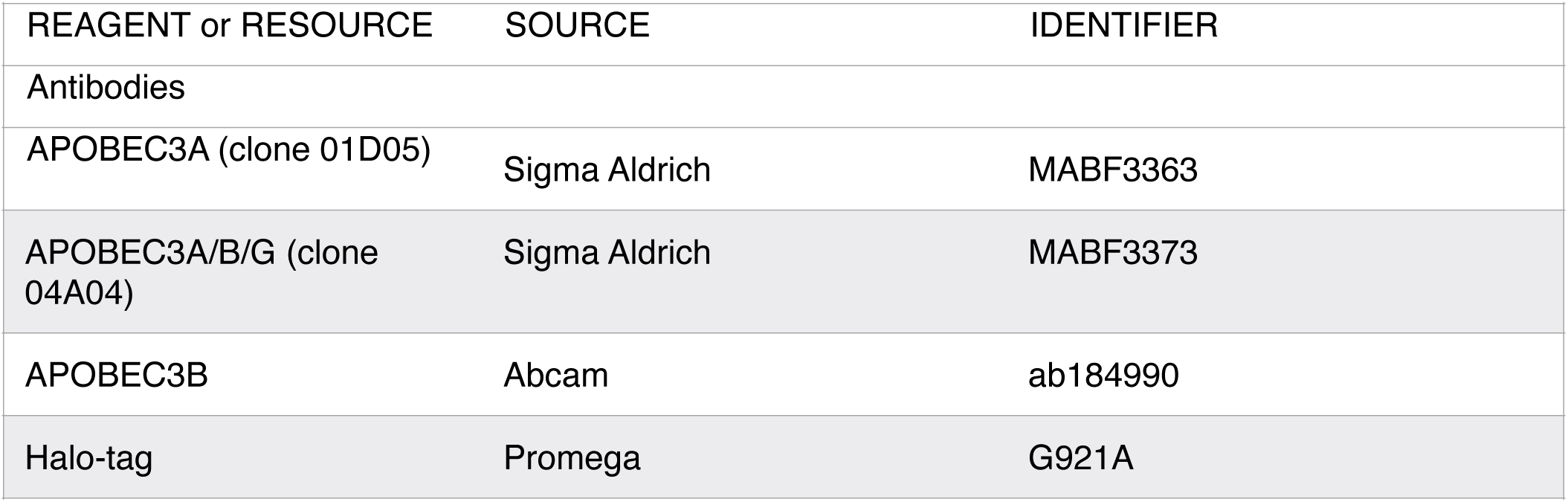
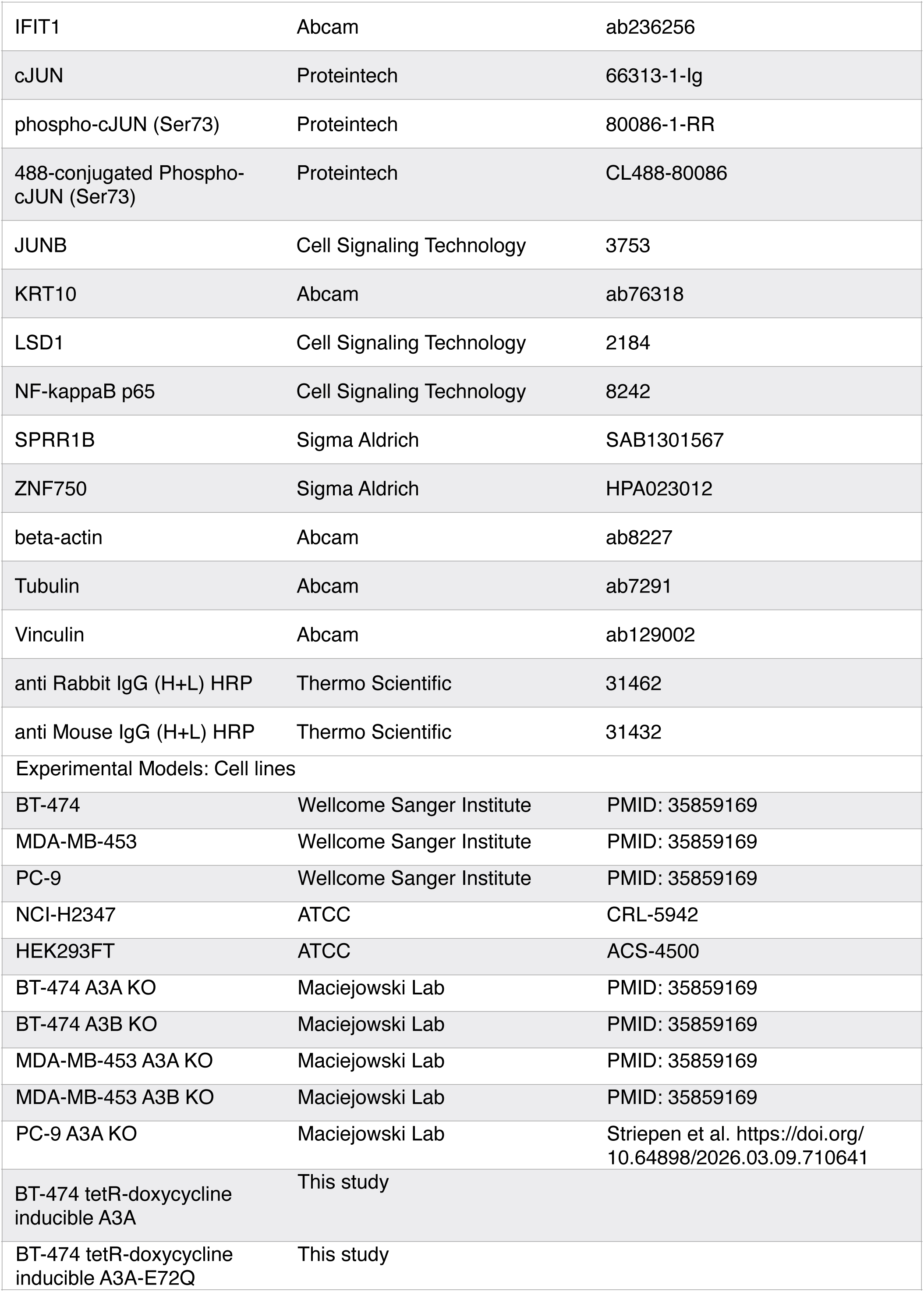
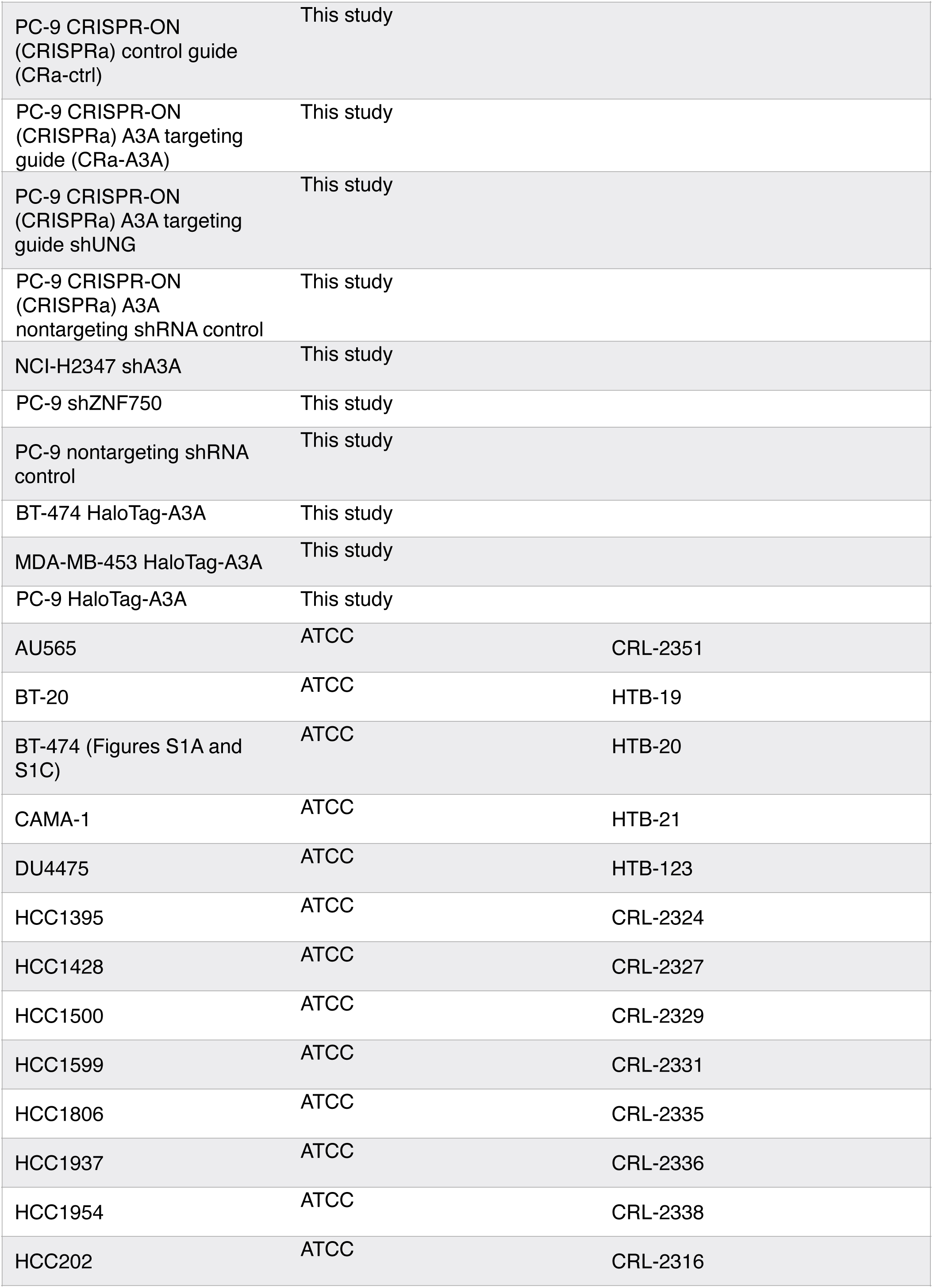
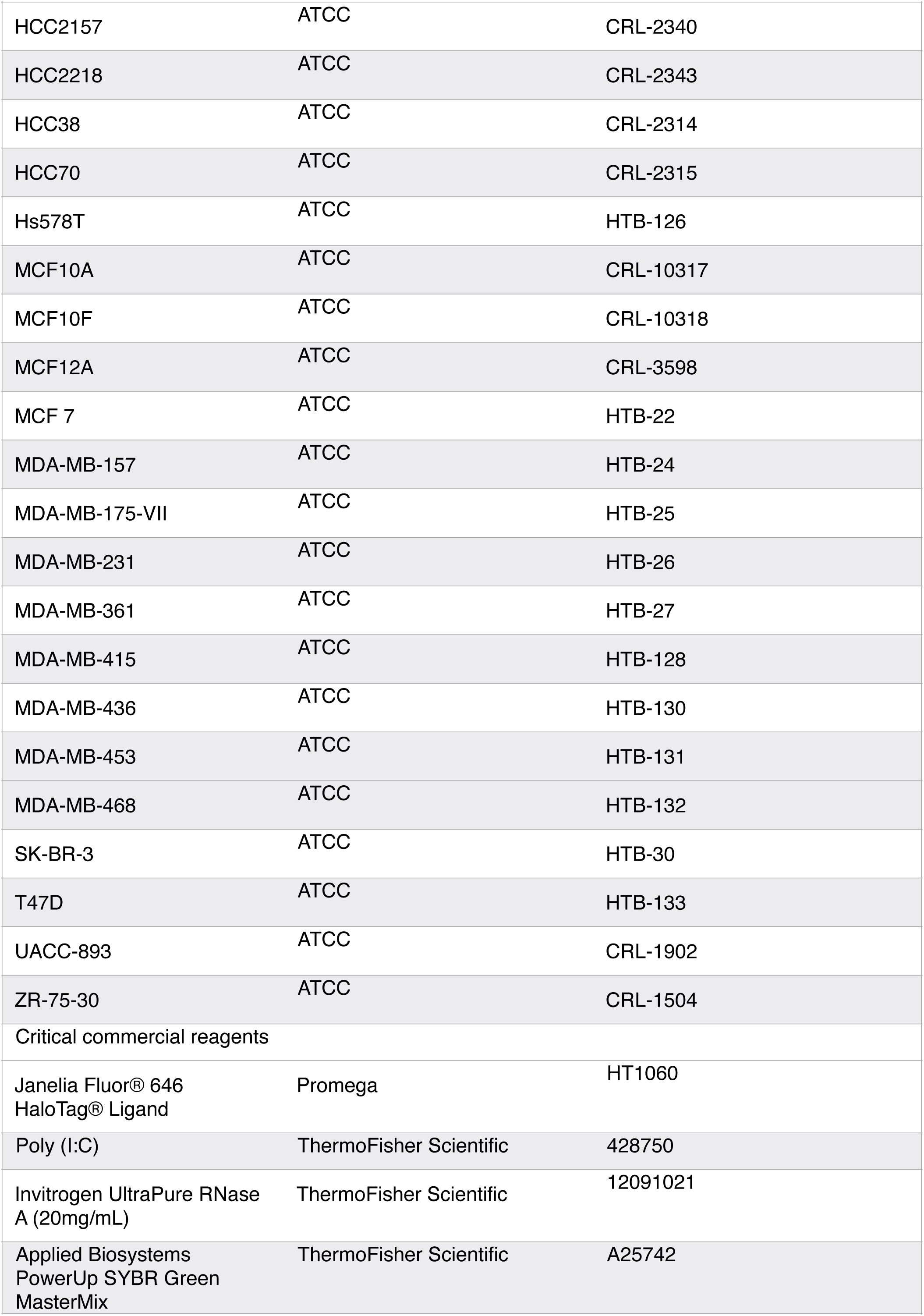
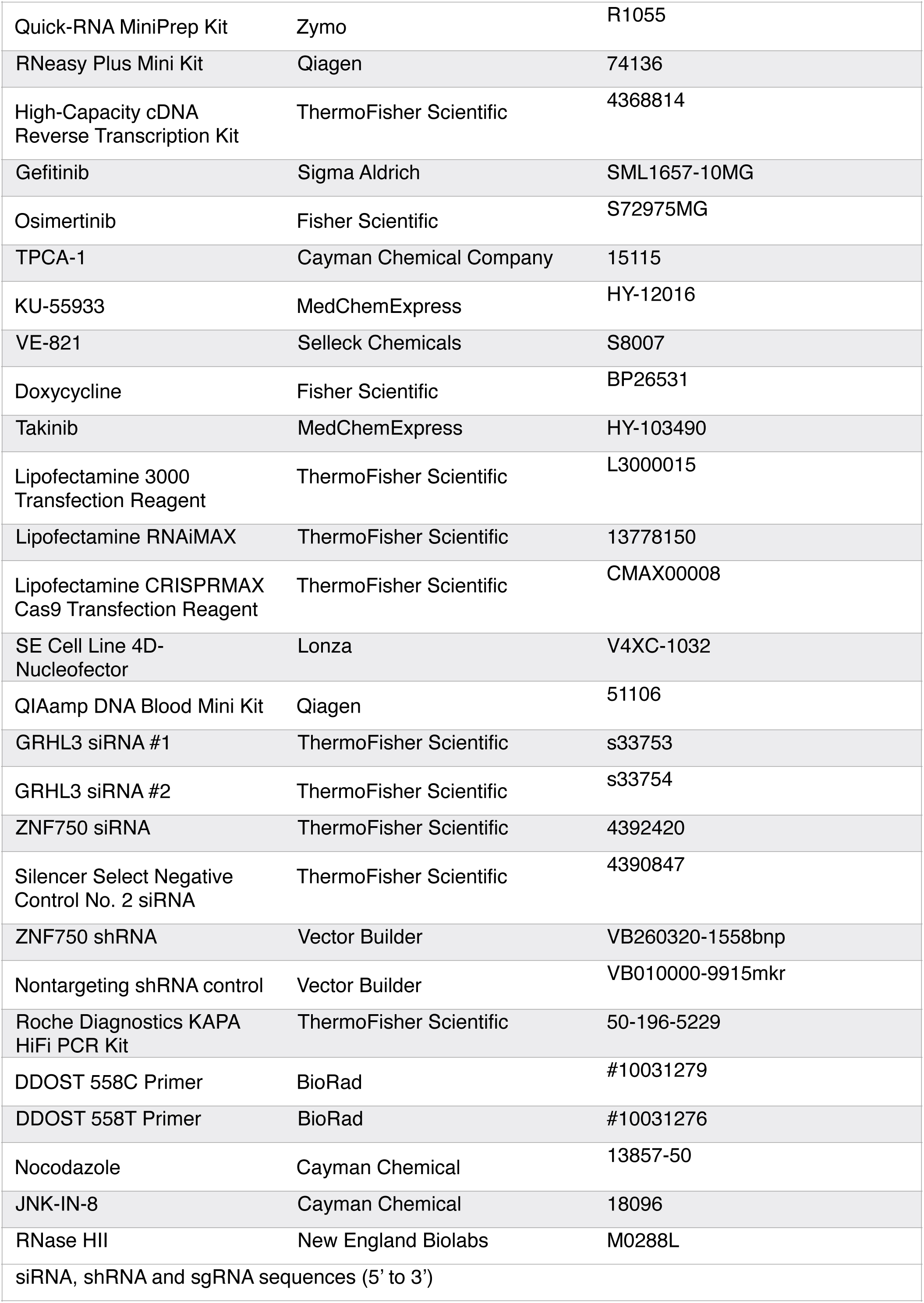
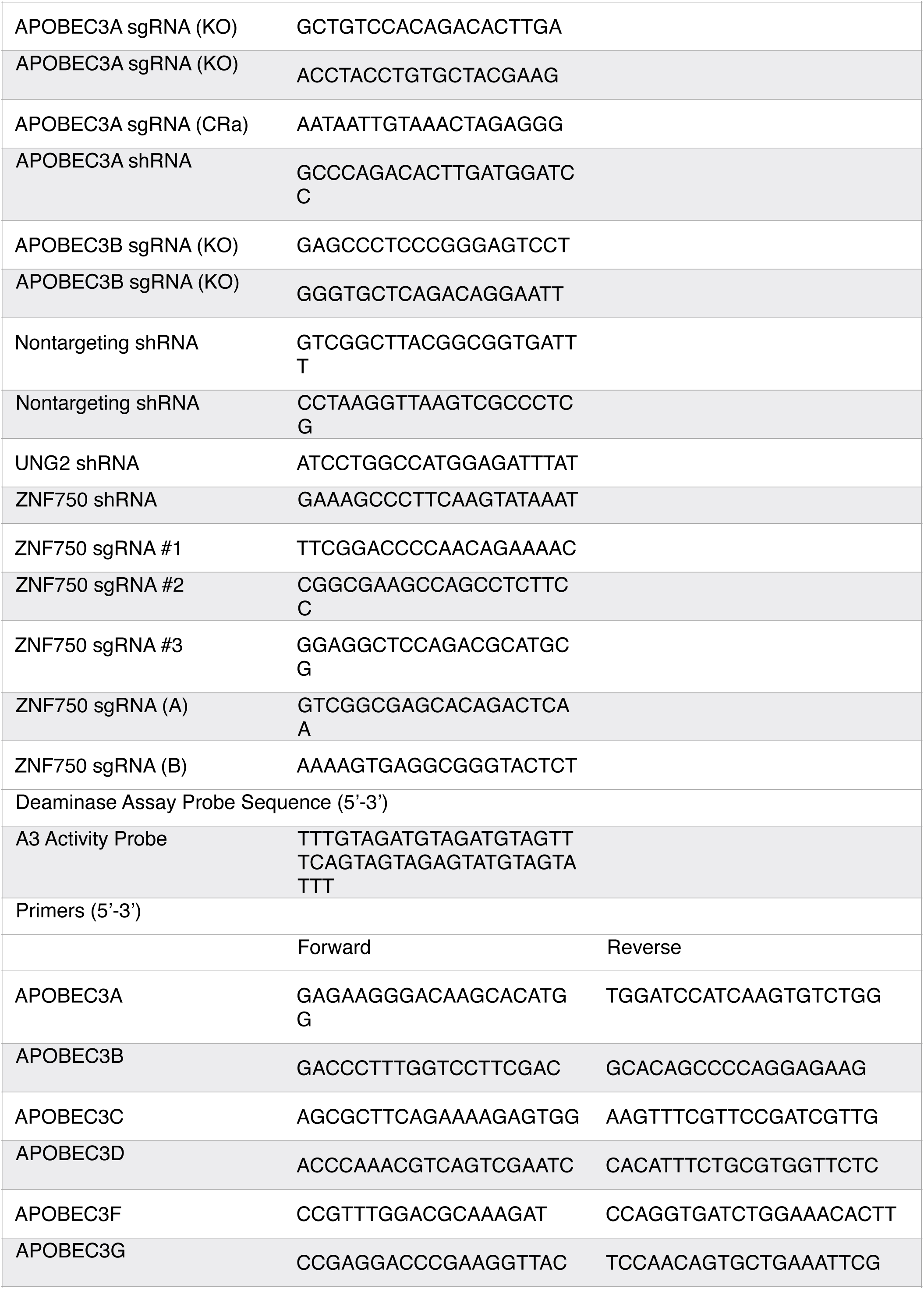
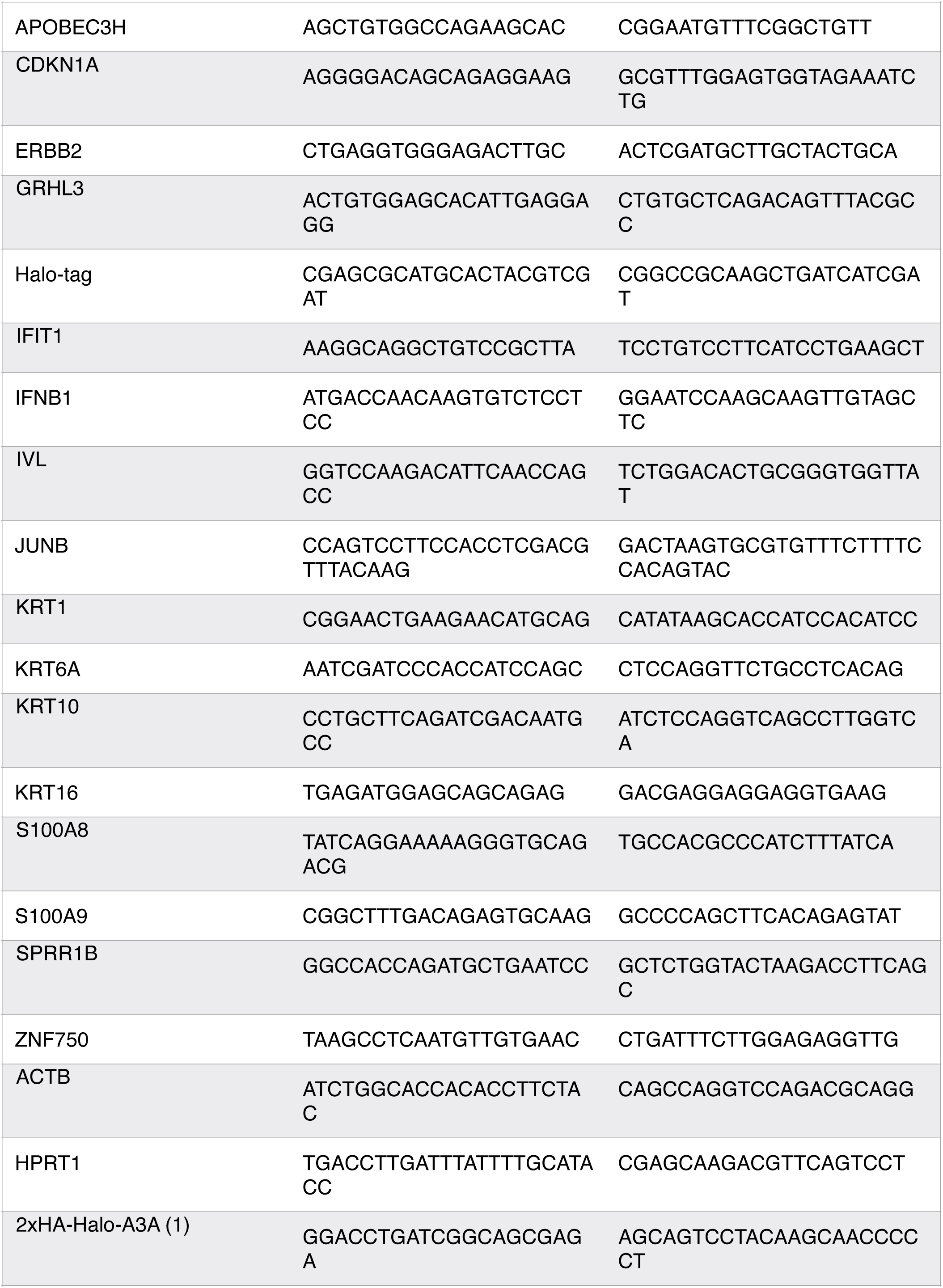
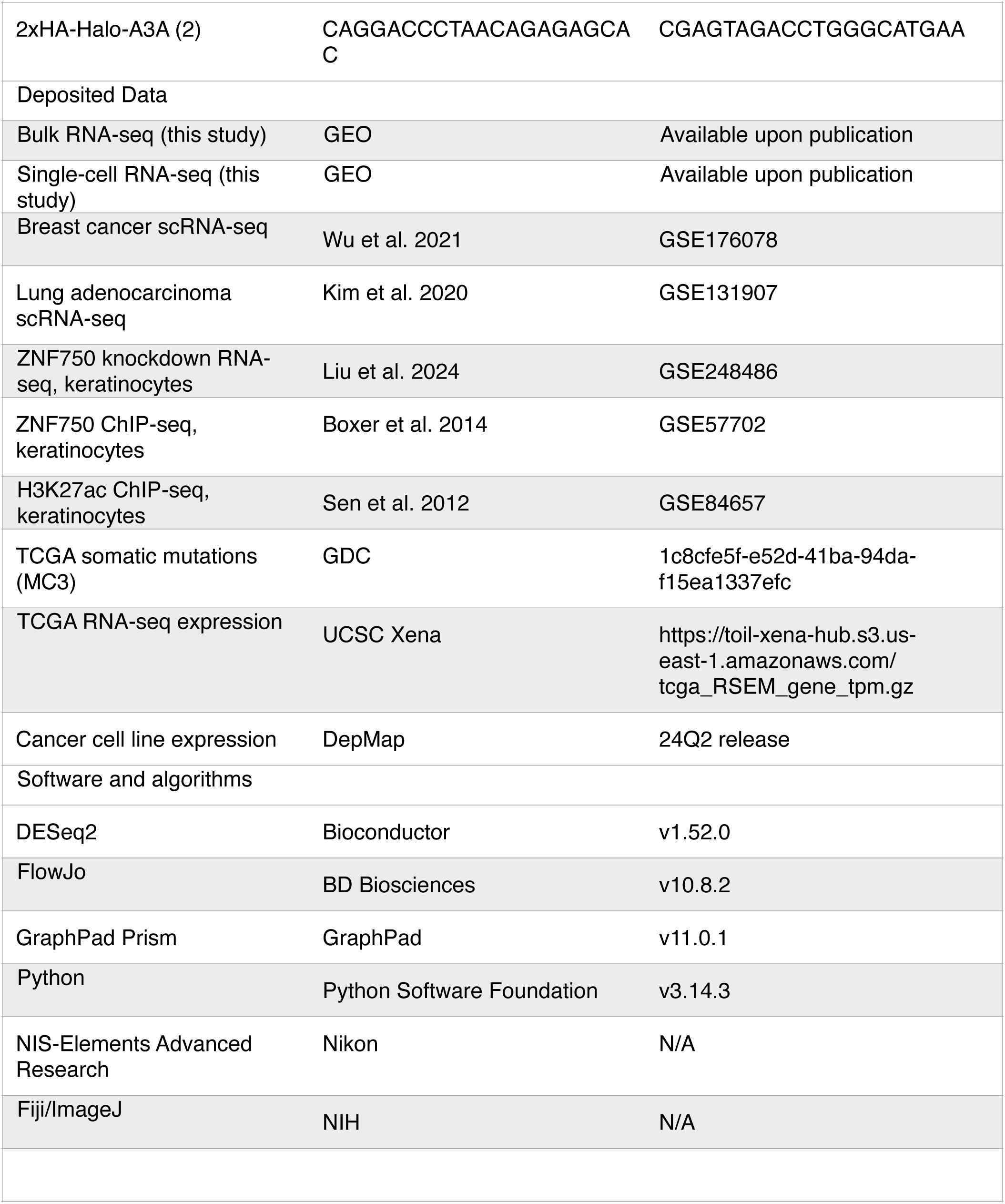

## RESOURCE AVAILABILITY

## Lead contact

Further information and requests for resources and reagents should be directed to and will be fulfilled by the Lead Contact, John Maciejowski.

## Materials availability

Cell lines generated in this study and listed in the Reagents and Resources Table are available from Dr. John Maciejowski.

## Data and code availability

RNA sequencing data have been deposited at GEO and will be made publicly available upon peer-reviewed publication. Previously published scRNA-seq data were obtained from GEO (Wu et al., GSE176078; Kim et al., GSE131907). ZNF750 knockdown RNA-seq data were from Liu et al. (GSE248486). TCGA somatic mutation calls were obtained from the GDC MC3 Public MAF. TCGA RNA-seq expression data were obtained from the UCSC Xena Toil RNA-seq Recompute hub. Cancer cell line expression and dependency data were obtained from the DepMap portal (24Q2 release). ZNF750 transcription factor perturbation signatures were accessed via the Enrichr TF_Perturbations_Followed_by_Expression library, which includes data from CREEDS (GSE38039). Gene set enrichment analyses used MSigDB v6.0 and MSigDB 2026.1.Hs gene set collections.

All custom analysis code and figure-generation scripts used in this study will be made publicly available via GitHub upon peer-reviewed publication. The repository contains Python and R scripts for RNA-seq differential expression visualization, gene set enrichment analysis, single-cell RNA-seq reanalysis, TCGA pan-cancer analyses, and APOBEC mutational signature quantification. The repository will be made publicly available upon acceptance of the manuscript.

## EXPERIMENTAL MODEL DETAILS

### Cell Culture

BT-474, MDA-MB-453 and PC-9 cell lines were acquired from collaborators or public repositories and extensively characterized as part of the Genomics of Drug Sensitivity in Cancer (GDSC) and COSMIC Cell Line projects^11–13,89^. NCI-H2347 cells were acquired from ATCC. Individual cell lines and derived clones were genotyped to confirm their identities^11^. All cell lines were routinely confirmed to be mycoplasma negative (LookOut Mycoplasma PCR Detection Kit, Sigma Aldrich MP0035-1KT). BT-474 and PC-9 cells were grown in RPMI medium supplemented with 10% fetal bovine serum (FBS), 1% penicillin-streptomycin (100 U/mL), 1% sodium pyruvate (1 mM), and 1% glucose (2 mg/mL). MDA-MB-453 cells were grown in DMEM:F12 medium supplemented with 10% FBS and 1% penicillin-streptomycin (100 U/mL). NCI-H2347 cells were grown in RPMI medium supplemented with 10% FBS and 1% penicillin-streptomycin (100 U/mL). Unless otherwise noted, all media and supplements were supplied by the MSKCC Media Preparation core facility. For pan-cell line analysis of *APOBEC3A* and *APOBEC3B* expression, AU565, BT20,BT-474, Cama1, DU4475, HCC1395, HCC1428, HCC1500, HCC1599, HCC1806, HCC1937, HCC1954, HCC202, HCC2157, HCC2218, HCC38, HCC70, Hs578T, MCF10A, MCF10F, MCF12A, MCF7, MDA-MB-157, MDA-MB-175-VII, MDA-MB-231, MDA-MB-361, MDA-MB-415, MDA-MB-436, MDA-MB-453, MDA-MB-468, SKBR3, T47D, UACC893, and Zr-75-30 cells were obtained from ATCC and cultured according to ATCC recommendations.

Poly(I:C) (200 ng/µL) was transfected using Lipofectamine 3000 (Invitrogen, L3000015) 24 hours before flow cytometry. Gefitinib (1 µM) and osimertinib (1 µM) were applied for 48 hours. Doxycycline was used at 1 µg/mL for inducible constructs, with duration varying by experiment as specified in figure legends. JNK inhibitor (JNK-IN-8, Cayman Chemical, 18096) was used at 10 µM. TAK1 inhibitor (Takinib, MedChemExpress, HY-103490) was used at 10 µM. ATM inhibitor (KU-55933, MedChemExpress, HY-12016) was used at 10 µM and ATR inhibitor (VE-821, Selleck Chemicals, S8007) was used at *5* µM.

### Endogenous HaloTag-APOBEC3A Knock-in

A 2xHA-HaloTag cassette was inserted at the N-terminus of *APOBEC3A* exon 1 on chromosome 22q13.1 by CRISPR-mediated homology-directed repair. A donor plasmid was constructed in a pUC19 backbone using a gene block containing left and right homology arms (800 bp each) flanking the 2xHA-HaloTag sequence (IDT). pUC19 was linearized with Eco53kI and assembled by Gibson assembly (NEB). The assembled construct was verified by Sanger sequencing. One million cells (BT-474, MDA-MB-453, or PC-9) were transfected with the donor plasmid, guide RNA oligonucleotide (target sequence: ACCGAGAAGGGACAAGCACA), and Cas9 protein using Lipofectamine CRISPRMAX Cas9 Transfection kit (Thermo Fisher). One week post-transfection, cells were labeled with 200 nM JF646-HaloTag ligand (Janelia Research Campus), and HaloTag-positive cells were bulk sorted and subcloned by limiting dilution in 96-well plates using FACSAria (BD Biosciences). Clones were screened by PCR using Phusion High-Fidelity DNA Polymerase (Thermo Scientific) with two primer pairs: AD382/AD387 (within tag; F: GGACCTGA TCGGCAGCGAGA, R: AGCAGTCCTACAAGCAACCCCCT) and AD 388 / AD 391 (flanking insertion site; F: CAGGACCCTAACAGAGAGCAC, R: CGAGTAGACCTGGGCATGAA).

### Generation of Knockout Cell Lines

*APOBEC3A* and *APOBEC3B* knockout lines in BT-474 and MDA-MB-453 were generated as described previously^11^. PC-9 *APOBEC3A* knockout clones and wild-type subclones were generated as described^13^. Briefly, cells (2 × 10^6 per 10 cm plate) were transfected using Lipofectamine 3000 (Invitrogen, L3000015) with 10 µg of pU6-sgRNA_CBh-Cas9-T2A-mCherry plasmid DNA. Twenty-four hours post-transfection, mCherry-positive cells were single-cell sorted by FACS using FACSAria (BD Biosciences) into 96-well plates. Guide RNA sequences are listed in Table S6.

### Doxycycline-Inducible APOBEC3A

BT-474 cells were engineered with doxycycline-inducible wild-type APOBEC3A (iA3A) and catalytic-dead E72Q mutant (i-dA3A) constructs as described^90^. Briefly, cells were transduced with lentivirus constructed with pLenti_CMV_TetR_Blast (Addgene, Cat #17492) and selected with blasticidin to produce a population of BT-474 cells expressing TET-R. Subsequent transduction with inducible expression vectors and selection with puromycin was used to produce iA3A and i-dA3A cells, respectively. The inducible vectors were derived by modifications of pLenti CMV Puro DEST (Addgene, #17452) that reduced CMV promoter activity, introduced gateway cloning compatibility, and changed the orientation of the expression cartridge to enable production of high titer lentivirus. The E72Q catalytic-dead mutant was generated by site-directed mutagenesis. Details of vector construction are previously published^91^. For induction, parental BT-474 cells or iA3A cells were cultured to 60% confluence after which doxycycline was added to the media at a final concentration of 1 µg/mL. Cells were harvested 96 hours later for RNA extraction.

### CRISPRa System

A doxycycline-inducible dCas9-VPR construct (Addgene, Plasmid #63800) was delivered to PC-9 cells with a transposase using Lipofectamine 3000 (Invitrogen, L3000015) following the manufacturer’s recommendations. Cells were selected with 200 µg/mL Hygromycin B (Thermo Fisher, 10687010). Single-cell clones were derived by limiting dilution and screened for system induction by immunoblotting. Clones were then transduced with lentiviral constructs stably expressing a single-guide RNA targeting the *APOBEC3A* promoter (A3A sg) or a control sgRNA (Ctrl sg). Guide RNA plasmids were constructed using a pLenti-sgRNA backbone. Transduced cells were selected with 800 ng/mL puromycin (Thermo Fisher, A1113803). sgRNA sequences are listed in Table S6.

### shRNA and siRNA-Mediated Depletion

For *APOBEC3A* knockdown in NCI-H2347, and UNG2 and *ZNF750* knockdown in PC-9, cells were transduced with lentiviral shRNA targeting *APOBEC3A* (shA3A), *UNG2* (shUNG), or *ZNF750* (shZNF750), or a nontargeting control. Lentivirus was produced by co-transfecting HEK293FT cells with the shRNA plasmid, psPAX2, and pMD2.G (Addgene) using calcium phosphate precipitation. Supernatants were filtered and supplemented with 4 µg/mL polybrene. Transduced cells were selected with hygromycin B (shA3A: 200 µg/mL), by flow cytometry sorting on BFP-positive cells (shUNG), or with neomycin (shZNF750: 400 µg/mL).

For *ZNF750* and *GRHL3* depletion, BT-474 and PC-9 HaloTag-A3A cells were transfected with siRNA targeting ZNF750 (Thermo Fisher, 4392420), GRHL3 (Thermo Fisher, s33753 and s33754), or a nontargeting control (Thermo Fisher, 4390847) using Lipofectamine RNAiMAX Transfection Reagent (Thermo Fisher, 13778100). siRNA and shRNA target sequences are listed in Table S6.

### sgRNA-Mediated Depletion

For ZNF750 depletion by CRISPR-Cas9, PC-9, BT-474, MDA-MB-453, and NCI-H2347 cells were electroporated using the Lonza 4D-Nucleofector X Unit (program EW-113). Alt-R S.p. Cas9 Nuclease V3 (IDT, 1081058) was incubated with guide RNAs (Table S6) for 20 minutes and then electroporated into cells using the SE Cell Line 4D-Nucleofector X Kit (Lonza, V4XC-1032) per the manufacturer’s instructions. Reactions were supplemented with 25 units of RNase HII (NEB, M0288L) and 100 ng/mL nocodazole (Cayman, 13857-50).

### Flow Cytometry and FACS Sorting

HaloTag-A3A and parental cells were plated at approximately 70% confluency. Culture medium was replaced with warm medium containing 200 nM JF646-HaloTag ligand (Janelia Research Campus), and cells were incubated for 20 minutes at 37°C. Cells were washed three times with PBS, returned to full medium with drug replenished if applicable, and incubated for 4-6 hours before trypsinization for analysis or sorting. For PC-9 experiments, cells were treated with 1 µM gefitinib or 1 µM osimertinib for 48 hours prior to HaloTag labeling. Flow cytometry analysis was performed on a BD LSR Fortessa using BD FACSDiva software. Cell sorting was performed on a BD FACSAria or a Sony biotechnology SH800 high-speed cell sorter at the MSKCC Flow Cytometry Core Facility. Parental non-HaloTag cell lines labeled with JF646 under identical conditions were used as negative controls. Data were analyzed in FlowJo 10.8.2. DAPI (1 µg/mL) was added immediately prior to analysis to exclude dead cells by gating on DAPI-negative events. HaloTag-A3A-high cells were defined by gating relative to the parental control. HaloTag-A3A-low cells were defined as the lower quartile of the HaloTag-negative population.

For burst dynamics experiments, BT-474 HaloTag-A3A cells were sorted for the HaloTag-high fraction, returned to culture, and re-analyzed at 0, 48, 96, and 192 hours post-sort (n = 3 independent experiments), or returned to culture and re-analyzed at 192 hours post-sort (n=3 independent experiments).

For phospho-c-Jun (Ser73) flow cytometry, BT-474 HaloTag-A3A cells were treated with 200 nM JF646-HaloTag ligand and incubated for 20 minutes at 37°C. Cells were washed three times with PBS and returned to full medium for 4 hours. Cells were then collected and fixed in cold fixation buffer (2% PFA) at 1 × 10⁶ cells per 100 µL. Cells were permeabilized for 15 minutes in a permeabilization buffer (0.2% NP-40, 3% goat serum). Then 0.5 µg of conjugated phospho-c-Jun antibody was added per 1 × 10⁶ cells and cells were incubated for 30 minutes at 4°C in the dark. Cells were washed once with a 1.3 mL permeabilization buffer and analyzed by flow cytometry.

### Immunoblotting

Cells were lysed in RIPA buffer (150 mM NaCl, 50 mM Tris-HCl pH 8.0, 1% NP-40, 0.5% sodium deoxycholate, 0.1% SDS, Pierce Protease Inhibitor Tablet EDTA-free) or sample buffer (62.5 mM Tris-HCl pH 6.8, 0.5 M β-mercaptoethanol, 2% SDS, 10% glycerol, 0.01% bromophenol blue). Quantification of RIPA extracts was performed using the Pierce BCA Protein Assay kit (Thermo Fisher Scientific). Proteins were separated on 12% or 8-16% gradient SDS-PAGE gels and transferred by wet transfer using 1× Towbin buffer (25 mM Tris, 192 mM glycine, 0.01% SDS, 20% methanol) onto 0.45 µm nitrocellulose membrane. Blocking was performed in 5% milk in 1× TBST (19 mM Tris, 137 mM NaCl, 2.7 mM KCl, 0.1% Tween-20) for 1 hour at room temperature.

The following primary antibodies were diluted in 1% milk in 1× TBST: anti-APOBEC3A/B/G (04A04, Millipore MABF3373, 1:1,000)^11^, anti-APOBEC3A (01D05, Millipore Sigma MABF3363, 1:1,000)^11^, anti-APOBEC3B (Abcam, ab184990, 1:500), anti-HaloTag (Promega, G921A, 1:1,000), anti-IFIT1 (Abcam, ab236256, 1:1,000), anti-LSD1 (Cell Signaling Technology, 2184, 1:1,000), anti-β-actin (Abcam, ab8227, 1:1,000), anti-α-tubulin (Abcam, ab7291), anti-NF-κB p65 (Cell Signaling Technology, 8242, 1:1,000), anti-vinculin (Abcam, ab129002), anti-phospho-c-Jun Ser73 (Proteintech, 80086-1-RR, 1:1,000), anti-JUNB (Cell Signaling Technology, 3753, 1:1,000), anti-c-Jun (Proteintech, 66313-1-Ig, 1:1,000), anti-KRT10 (Abcam, ab76318, 1:1,000), anti-SPRR1B (Sigma Aldrich, SAB1301567, 1:500), anti-ZNF750 (Sigma-Aldrich, HPA023012, 1:500). Secondary antibodies: anti-mouse IgG HRP (Thermo Fisher Scientific, 31432, 1:10,000) and anti-rabbit IgG HRP (Thermo Fisher Scientific, 31462, 1:10,000). Generation and validation of monoclonal antibodies 04A04 and 01D05 were described previously^11^. A complete list of antibodies is provided in Table S7.

### Quantitative RT-PCR

RNA was isolated using the Quick-RNA MiniPrep Kit (Zymo Research, R1055) or the RNeasy Plus Mini Kit (Qiagen, 74136). cDNA was synthesized using High-Capacity cDNA Reverse Transcription Kit (Applied Biosystems, 4368814). cDNA synthesis reactions were performed as described in the manufacturer’s protocol. Quantitative PCR was performed on a QuantStudio 6 Pro Real-Time PCR system (Applied Biosystems) using PowerUp SYBR Green master mix (Applied Biosystems, A25742) / UPL probes (Roche). Expression was normalized to *β-actin* and *HPRT1* reference genes using the ΔΔCt method. For comparison of *APOBEC3A* and *APOBEC3B* mRNA abundance across a panel of breast cancer cell lines, RNA was extracted from cells during log-phase growth using the Omega Bio-Tek Total RNA kit. RNA was DNase treated with DNaseI (NEB), and subsequently combined with cDNA reaction mixture (2.5 μM oligo dT23VN and 3.5 μM random hexamers, 0.5 mM dNTPs, 1x Mashup RT Reaction Buffer (25mM Tris-HCl pH 8.3, 25 mM MOPS pH 7.9, 60 mM KCl, 4 mM MgCl^2^, 5% glycerol, 0.006% IGEPAL CA-630), 10 mM DTT, 0.5 μL Mashup Reverse Transcriptase and 16 U RNase Inhibitor (Ribolock Rnase Inhibitor, ThermoFisher). Reactions were then incubated at 42°C for 1 hr, then 65°C for 20min. qPCR was performed using 2 µL cDNA with Forget-Me-Not EvaGreen qPCR master mix (Biotium) and 2 µM primers for a total reaction volume of 20 µL using a BioRad CFX96. Cycling parameters were as follows: 95°C for 5 mins, then 40 cycles of 95°C for 10s and 62.5°C for 1 min. *APOBEC3A* and *APOBEC3B* mRNA abundances were determined relative to *HPRT1* mRNA. Primer sequences are listed in Table S8.

### Subcellular Fractionation

BT-474 and MDA-MB-453 cells were fractionated into cytoplasmic and nuclear compartments using the REAP method. Briefly, 2 million cells were washed with ice-cold PBS, scraped, and centrifuged for 10 seconds. The pellet was resuspended in 900 µL ice-cold 0.1% NP-40 in PBS and an aliquot was removed as whole-cell lysate. After centrifugation for 10 seconds, the supernatant was collected as the cytosolic fraction and combined with 1× Laemmli sample buffer. The nuclear pellet was resuspended in 1 mL ice-cold 0.1% NP-40 in PBS, centrifuged again, and the final pellet was resuspended in 1× Laemmli sample buffer. Nuclear fractions and whole-cell lysates were sonicated and boiled for 1 minute at 95°C. LSD1 served as a nuclear marker and tubulin as a cytoplasmic marker. Representative of n = 2 independent experiments.

### Live-Cell Fluorescence Microscopy

FACS-sorted HaloTag-A3A-positive BT-474 cells were plated on poly-L-lysine-coated 4-chamber 35 mm glass-bottom dishes and incubated with JF646-HaloTag ligand 6 hours before imaging. Cells stably expressing H2B-mCherry (Addgene, 164244)^92^ were used for chromatin colocalization. Images were acquired on a Nikon SoRa Spinning Disk Confocal system with Borealis microadapter, Perfect Focus 4, motorized turret and encoded stage, PRIME 95B Monochrome Digital Camera, and a CFI Apo TIRF 60x 1.49 NA objective lens (W.D. 0.12mm). NIS-Elements Advanced Research Software was used for acquisition on a Dual Xeon Imaging workstation. Images were denoised using default parameters. Maximum intensity projects and brightness/contrast adjustments were performed in Fiji. Scale bars, 10 µm.

### Clonogenic Survival and Proliferation Assays

For clonogenic assays, 1,000 sorted cells were plated in duplicate in 6-well plates and cultured for 18 days. Cells were fixed with methanol, rinsed with PBS, and stained with 0.4% (w/v) crystal violet in 20% methanol for 5 minutes. Colonies were counted using the “Colony Area” plugin in Fiji ImageJ and plotted as clonogenic survival relative to parental cells.

For proliferation assays, cells were seeded in triplicate in 96-well plates and confluence was monitored by daily imaging on an IncuCyte Live-Cell Analysis Imager (Essen/Sartorius). Technical replicates were averaged and plotted over time to generate growth curves.

### *In Vitro* DNA Deaminase Activity Assay

Deamination activity assays were performed as described^11,13^. Briefly, cells (∼10^6^) were pelleted and lysed in deamination lysis buffer (25 mM HEPES pH 8.0, 150 mM NaCl, 1 mM EDTA, 10% glycerol, 0.5% Triton X-100, protease inhibitor cocktail). Lysates were sheared through a 28.5-gauge syringe and cleared by centrifugation at 13,000 × g for 20 minutes at 4°C. Protein was quantified by BCA assay and 50 µg of protein was used per reaction. Reactions were assembled in 50 µL containing lysate, 1× deaminase buffer (225 mM Tris-HCl pH 7.5, 20 mM KCl, 25 mM NaCl, 0.05% Triton X-100, 10 mM DTT), 0 . 1 µ M IRDye 800 - labeled TC-containing oligonucleotide probe (5 ′- IRD 800 / T*T*T*GTAGATGTAGATGTAGTTTCAGTAGTAGAGTATGTAGT*A*T*T*T), 1.5 units UNG (NEB, M0372L), and ±0.5 µL RNase A (20 mg/mL, Thermo Fisher, 12091021). Reactions were incubated at 37°C with aliquots removed at indicated time points. Each aliquot was treated with 2 µL of 2 M NaOH and incubated at 95°C for 12 minutes to cleave abasic sites, then neutralized with 2 µL of 2 M HCl. Reactions were terminated by adding an equal volume of Novex TBE-Urea Sample Buffer 2× (Thermo Fisher, LC6876) and separated on a 15% TBE-Urea gel (Thermo Fisher, EC68852BOX) in 1× TBE buffer for 120 minutes at 160 V. Gels were imaged on an Odyssey Infrared Imaging System (LI-COR) and quantified using ImageJ. Percent deamination was fit to a one-site binding hyperbola in GraphPad Prism Version 11.0.1.

### *DDOST* RNA Editing Assay

*DDOST* 558C>U RNA editing assays were performed as described previously^11,29^ with assistance from the MSKCC Integrated Genomics Operation. Total RNA was extracted using the RNeasy Plus Mini kit (Qiagen, 74136). cDNA was generated using the High Capacity cDNA Reverse Transcription Kit (Thermo Fisher Scientific, 4368814). cDNA (20 ng) along with primers (Bio-Rad, 10031279 and 10031276) for the target *DDOST*^558C>U^ amplification was mixed in PCR reactions in a total volume of 25 µL. Twenty microliters of the reaction was mixed with 70 µL of Droplet Generation Oil for Probes (Bio-Rad) and loaded into a DG8 cartridge (Bio-Rad). A QX200 Droplet Generator (Bio-Rad) was used to generate droplets, which were transferred to a 96-well plate. PCR cycling: 5 minutes at 95°C, 40 cycles of 94°C for 30 seconds and 53°C for 1 minute, and 98°C for 10 minutes. The QX200 Droplet Reader (Bio-Rad) was used for fluorescence measurement of FAM and HEX probes. Data were analyzed using QuantaSoft software (Bio-Rad) with gating based on positive and negative DNA oligonucleotide controls.

### Single-Cell RNA Sequencing

BT-474 and MDA-MB-453 single-cell RNA sequencing was performed using a plate-based Smart-seq2-like protocol. Single cells were sorted into Buffer RLT (Qiagen, 79216) and RNA was purified using Agencourt RNAClean XP beads (Beckman Coulter, A63987) at a 2.2× ratio. First-strand cDNA synthesis was performed using Maxima H Minus Reverse Transcriptase (ThermoFisher, EP0751) with oligo dT primers and a custom template-switch oligo at 1 mM final concentration. cDNA was amplified for 24 cycles using KAPA HiFi HotStart ReadyMix (Kapa Biosystems, KK2601). After PicoGreen quantification, 0.2 ng of cDNA was used to prepare libraries with the Nextera XT DNA Library Preparation Kit (Illumina, FC-131-1024) in a total volume of 6.25 µL with 8-12 cycles of PCR. Indexed libraries were pooled by volume and cleaned with AMPure XP beads (Beckman Coulter, A63882) at a 0.6-2× ratio. Pools were sequenced on a NovaSeq 6000 in a PE150 run using the NovaSeq 6000 S1 Reagent Kit (300 cycles, Illumina). An average of 4.6 million paired reads were generated per sample and the percent of mRNA bases per sample averaged 48%. Gene-level counts were quantified using HTSeq. Quality control filtering retained cells with at least 100 detected genes and no more than the 99th percentile of genes per cell. Genes detected in fewer than 2 cells were excluded.

Data were processed using scanpy 1.12. Counts were normalized per cell to a target sum of 10,000 and log1p-transformed. The top 2,000 highly variable genes were selected using the Seurat flavor in scanpy. All seven APOBEC3 paralogs (APOBEC3A, APOBEC3B, APOBEC3C, APOBEC3D, APOBEC3F, APOBEC3G, APOBEC3H) were excluded from the highly variable gene set to prevent APOBEC3A expression from driving dimensionality reduction structure. APOBEC3H was not detected at the minimum threshold of 2 cells and was filtered during preprocessing. PCA was performed with 30 components. A neighbor graph was constructed with n_neighbors = 15 and n_pcs = 30. UMAP was computed with min_dist = 0.3 and random_state = 42. All computations were performed jointly on both cell lines. APOBEC3A detection was defined as raw count ≥ 1.

### Bulk RNA Sequencing and Differential Expression

For BT-474 Halo-sort and PC-9 osimertinib experiments, after RiboGreen quantification and quality control by Agilent BioAnalyzer, 109-500 ng of total RNA with RIN values of 8.9-10 underwent polyA selection and library preparation using the TruSeq Stranded mRNA LT Kit (Illumina, RS-122-2102) according to the manufacturer’s instructions with 8 cycles of PCR. Samples were barcoded and run on a NovaSeq 6000 in a PE100 run using the NovaSeq 6000 S4 Reagent Kit (200 cycles, Illumina). An average of 15 million paired reads was generated per sample. Ribosomal reads represented 0.03-0.22% of the total reads generated and the percent of mRNA bases averaged 96%. For BT-474 doxycycline-inducible APOBEC3A experiments, RNA was isolated using the E.Z.N.A total RNA kit (Omega Bioscience), treated with DNase I, and further purified by phenol:chloroform/ethanol precipitation. RNA was submitted to Novogene (Sacramento, CA) for non-stranded eukaryotic mRNA-Seq on an Illumina sequencing platform generating 150 base paired-end reads.

Differential expression analysis was performed using DESeq2 (Wald test, Benjamini-Hochberg FDR correction). Log₂ fold change values are unshrunken maximum likelihood estimates. The following contrasts were analyzed: BT-474 HaloTag-A3A-high versus HaloTag-A3A-low (n = 2 per condition, 22,147 genes tested). BT-474 doxycycline-induced APOBEC3A versus uninduced controls (n = 2 per condition, 23,121 genes tested). PC-9 wild-type osimertinib versus untreated (n = 2 per condition, 19,393 genes tested). PC-9 APOBEC3A knockout osimertinib versus untreated (n = 2 per condition, 19,721 genes tested). PC-9 HaloTag-A3A-high versus HaloTag-A3A-low under osimertinib treatment (n = 2 per condition, 21,281 genes tested).

For principal component analysis, count matrices were filtered to genes with mean normalized counts > 1, log₂(x + 1) transformed, and standardized per gene. PCA was computed with n_components = 4, and the first two components were plotted.

### Gene Ontology and Gene Set Enrichment Analysis

Gene ontology over-representation analysis was performed using clusterProfiler [version ≥ 4.18] with org.Hs.eg.db [version ≥ 3.22] in R 4.5.2. The enrichGO function was applied to upregulated DEGs (padj < 0.05 and log∼2∼FC > 1) against a background of all Entrez-mapped genes with GO biological process annotation from the DESeq2 output. Redundant terms were collapsed using simplify (cutoff = 0.6, by = “p.adjust”). For the BT-474 Halo-sort analysis (Figure 2D), 17 upregulated DEGs were tested against a background of 12,888 Entrez-mapped genes.

Gene set enrichment analysis was performed using two methods. For the BT-474 Halo-sort dataset, GseaPreranked was run in GSEA v4.3.2 (Broad Institute) using the MSigDB v6.0 C5 GO biological process collection, the classic enrichment statistic, 1,000 gene-set permutations, and genes ranked by log∼2∼ fold change. For PC-9 datasets and BT-474 Dox-OE analyses, fgsea v1.36.2 was used in R 4.5.2 with MSigDB 2026.1.Hs C5 GO biological process gene sets accessed via msigdbr v26.1. Parameters: minSize = 10, maxSize = 500, nPermSimple = 10,000, seed = 42, ranked by DESeq2 Wald statistic.

For selected pathway enrichment panels, non-redundant terms were selected by greedy top-down deduplication, removing terms with leading-edge overlap exceeding 50% with an already-selected term. Where indicated, pathways were filtered to GO biological process terms with padj < 0.05 and NES > 0, deduplicated by leading-edge Jaccard filter (threshold 0.50), and up to three skin-related terms were force-included. Up to 15 terms by NES were displayed.

### Reanalysis of Public Datasets DepMap/CCLE

Bulk expression data were obtained from the DepMap portal (24Q2 release; https://depmap.org/portal/ download/all/). Gene expression values are reported as log∼2∼(TPM + 1) from the file OmicsExpressionTPMLogp1HumanProteinCodingGenes.csv. Cell line metadata were obtained from Model.csv. Lineage groupings for Figure 1A: breast (n = 70), bladder (n = 38), cervical (n = 22), head and neck (n = 75), lung adenocarcinoma (n = 79), lung squamous (n = 27), esophageal (n = 86), pooled APOBEC-high (n = 433). Kernel density estimation was performed with bandwidth = 0.18 over 400 points from 0 to 9.5.

### Wu et al. 2021 breast cancer scRNA-seq

Single-cell RNA sequencing data from 26 breast cancer patients were obtained from GEO (GSE176078)^27^. The dataset was filtered to cancer epithelial cells (celltype_major = “Cancer Epithelial”) with at least 200 detected genes per cell, yielding 24,489 cells. APOBEC3A detection rate: 0.7% (173/24,489 cells, raw count ≥ 1).

For correlation analyses, counts were normalized to 10,000 per cell and log1p-transformed. Genes detected in fewer than 5% of cells were excluded. Spearman correlations between *APOBEC3A* or *APOBEC3B* expression and all retained genes were computed, with Benjamini-Hochberg FDR correction. Hits were defined as ρ > 0.05 and FDR < 0.05.

### Kim et al. 2020 lung adenocarcinoma scRNA-seq

Single-cell RNA sequencing data were obtained from GEO (GSE131907)^57^. The dataset was filtered to primary tumor malignant epithelial cells (Sample_Origin = “tLung” with Cell_type = “Epithelial cells”, or Sample_Origin = “tL/B” with Cell_subtype = “Malignant cells”), with at least 200 detected genes per cell. Non-malignant, lymph node metastasis, brain metastasis, and pleural effusion samples were excluded. This yielded 13,670 cells from 15 patients. Normalization and correlation analysis were performed identically to the Wu et al. dataset. 10,737 genes were tested. *KRT16* was the top *APOBEC3A* correlate (ρ = 0.168, FDR = 6.3 × 10^−83). *GRHL1* ranked 13th among *APOBEC3A* correlates (ρ = 0.118, FDR = 1.8 × 10^−40).

### TCGA pan-cancer expression analysis

TCGA pan-cancer RSEM gene expression data were obtained from UCSC Xena (file: tcga_RSEM_gene_tpm.gz; pre-transformed as log∼2∼(TPM + 1)). Primary tumors were selected by sample barcode across all available cancer types, yielding 9,186 tumors across 32 TCGA cancer types. Pan-cancer correlations were computed across all 32 types. Ten cancer types with established APOBEC activity or squamous biology (BLCA, LUAD, LUSC, BRCA, HNSC, CESC, COAD, READ, ESCA, STAD) are highlighted in the scatter plot and used for the per-gene, per-cancer-type heatmap.

A 22-gene keratinocyte differentiation signature was defined by curating canonical squamous differentiation markers: *KRT5*, *KRT6A*, *KRT6B*, *KRT14*, *KRT16*, *KRT17*, *IVL*, *FLG*, *LOR*, *SPRR1A*, *SPRR1B*, *SPRR2A*, *SPRR2D*, *DSG1*, *DSG3*, *TGM1*, *TGM5*, *CNFN*, *EVPL*, *S100A7*, *S100A8*, *S100A9*.

*LOR* was not present in the UCSC Xena RSEM gene name mapping, yielding a 21-gene effective signature. The keratinocyte differentiation score for each tumor was computed as the mean of per-gene z-scores across all 9,186 samples. Genes were retained within each cohort if detected in >50% of samples. Pearson and Spearman correlations between *APOBEC3A* or *APOBEC3B* expression and the keratinocyte score were computed. The comparison of dependent correlations was performed using the Meng, Rosenthal, and Rubin (1992) test, accounting for the *APOBEC3A*-*APOBEC3B* inter-correlation across n = 9,186 tumors.

For per-gene, per-cancer-type Spearman correlations, 10 APOBEC-relevant cancer types were selected (BLCA, LUAD, LUSC, BRCA, HNSC, CESC, COAD, READ, ESCA, STAD). Individual gene × cancer-type pairs were corrected using Benjamini-Hochberg FDR.

### TCGA ZNF750 loss-of-function and APOBEC mutational burden

Somatic mutations were obtained from the GDC MC3 Public MAF (file UUID: 1c8cfe5f-e52d-41ba-94da-f15ea1337efc) via the GDC API. Per-sample APOBEC mutational burden (combined SBS2 + SBS13) was quantified using SigProfilerAssignment v1.1.3 with cosmic_fit (context_type = 96, genome_build = GRCh37, cosmic_version = 3.3). *ZNF750* loss-of-function was defined as Nonsense_Mutation, Frame_Shift_Del, Frame_Shift_Ins, Splice_Site, or Splice_Region mutations. Analysis was restricted to five squamous-enriched cancer types: HNSC (486 WT, 11 LOF), ESCA (171 WT, 9 LOF), CESC (277 WT, 9 LOF), BLCA (400 WT, 5 LOF), LUSC (473 WT, 6 LOF), pooled (1,807 WT, 40 LOF). Per-cancer-type significance was assessed by two-sided Mann-Whitney U test, with Benjamini-Hochberg adjustment across six tests. Pan-cancer analysis used OLS linear regression of log∼10∼(APOBEC burden + 1) on *ZNF750* LOF status with cancer-type dummy variables (coefficient = +0.93, p = 2.2 × 10^−11). Ninety-five percent bootstrap percentile confidence intervals were computed from 10,000 resamples.

### Liu et al. 2024 ZNF750 data

*APOBEC3* paralog expression in *ZNF750*-depleted versus control primary human keratinocytes was imported directly from^72^ Tables S1 and S2 (GEO: GSE248486). Linear fold-change values were obtained from the “Ratio (ZNF750i vs. CTLi)” column of Table S1. FDR values were from the “FDR step up” column. Direct ZNF750 ChIP targets were classified from Table S2 (ChIP-seq peak and knockdown-upregulated overlap). No local reanalysis of raw sequencing data was performed.

### Enrichr transcription factor perturbation analysis

The 17 upregulated DEGs from the BT-474 Halo-sort dataset (padj < 0.05, log∼2∼FC > 1) were submitted as input to the Enrichr API (https://maayanlab.cloud/Enrichr/) via gseapy v1.1.12, querying the TF_Perturbations_Followed_by_Expression library. The top enrichment hit was the response to *ZNF750* knockdown in primary human keratinocytes (CREEDS GSE38039). A curated ZNF750-regulated gene set was assembled as the union of Boxer et al. 2014 directly repressed targets (13 genes: *CDSN*, *DMKN*, *FLG*, *IVL*, *KLK7*, *KRT9*, *S100A7*, *SERPINB3*, *SPINK5*, *SPRR1A*, *SPRR1B*, *SPRR3*, *TMEM154*) and CREEDS GSE38039 ZNF750-KD upregulated genes (11 genes), yielding 15 unique genes after union^74^. Overlap between the 17 Halo-sort DEGs and this reference set was assessed by one-sided Fisher’s exact test (scipy.stats.fisher_exact, alternative = “greater”; OR = 681, p = 2.4 × 10^−10).

### Statistical Analyses

Statistical comparisons were performed using the tests indicated in individual figure legends. For pairwise comparisons, two-tailed Student’s *t*-test was used unless otherwise specified. For multi-group comparisons, one-way ANOVA with Tukey’s multiple-comparisons test was used. DESeq2 Wald test with Benjamini-Hochberg FDR correction was used for all RNA-seq differential expression analyses. P-value thresholds: *P < 0.05, **P < 0.01, ***P < 0.001, ****P < 0.0001, ns = not significant. Error bars represent mean ± s.d. unless otherwise noted. All wet-lab statistical analyses were performed in GraphPad Prism (Version 11.0.1).

### Data Reporting

No statistical methods were used to predetermine sample size. The investigators were not blinded to allocation during experiments and outcome assessment.

### Software

Python analyses used Python 3.14.3 with pandas 2.3.3, numpy 2.4.3, scipy 1.17.1, statsmodels 0.14.6, matplotlib 3.10.8, seaborn 0.13.2, scanpy 1.12, anndata 0.12.10, gseapy 1.1.12, SigProfilerAssignment 1.1.3, SigProfilerMatrixGenerator 1.3.6, scikit-learn 1.8.0, and umap-learn 0.5.11. R analyses used R 4.5.2 with fgsea 1.36.2, msigdbr 26.1, clusterProfiler [version ≥ 4.18], and org.Hs.eg.db [version ≥ 3.22]. DESeq2 [1.52.0]. Broad GSEA desktop application v4.3.2 was used for GseaPreranked analysis with MSigDB v6.0. Flow cytometry data were analyzed in FlowJo 10.8.2. Large language model tools were used for manuscript editing and consistency checking. All scientific content, data interpretation, and writing decisions were made by the authors.

## ACKNOWLEDGEMENTS

We thank members of the Maciejowski laboratory for reading the manuscript, M. Petljak for helpful discussions, and the MSKCC Flow Cytometry Core and Integrated Genomics Operation for technical support. We acknowledge the use of the Integrated Genomics Operation Core (RRID: SCR_027801), funded by the NCI Cancer Center Support Grant (CCSG, P30 CA008748). Work in J.M.’s laboratory is supported by the NCI (R01CA270102, R37CA261183, R01CA304441, and P30CA008748), the Pershing Square Sohn Cancer Research Alliance, and the Alan and Sandra Gerry Metastasis and Tumor Ecosystems Center. J.M. and S.A.R. are also supported by NCI R01CA269784. J.S. is supported by an NIH Ruth L. Kirschstein Predoctoral Individual National Research Service Award (F31CA290850). The Center for Epigenetics Research acknowledges funding by the NCI Cancer Center Support Grant (CCSG, P30 CA008748), Cycle for Survival, and the Metropoulos Foundation. S.A.R. acknowledges support from the NIH (R01CA269784).

## AUTHOR CONTRIBUTIONS

J.S., A.D., and J.M. conceived the study and designed experiments. J.S. and A.D. performed most experiments and data analysis, with technical support from A.R. R.K. performed RNA sequencing analysis. S.A.R. provided reagents, cell lines, RNA sequencing data, and contributed to characterization of the breast cancer cell line panel. E.T., A.H., A.N., H.R., C.C., T.M.M. and R.X.N. contributed to experiments and data analysis. J.M. supervised the study. J.S. and J.M. wrote the manuscript, with contributions from A.D. and input from all authors.

## SUPLEMENTARY FIGURES

**Figure S1.**
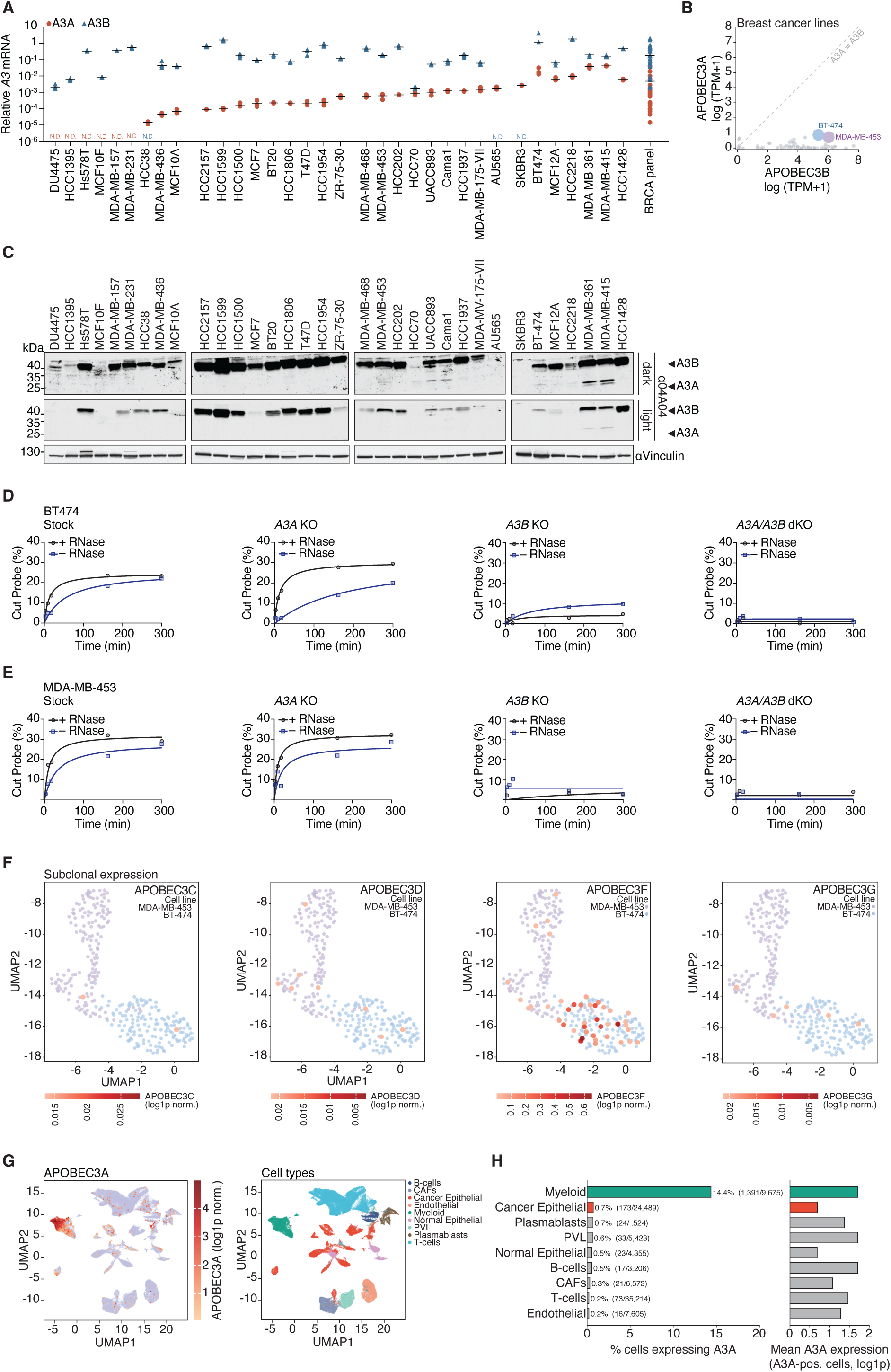
Bulk APOBEC3A expression and activity across cancer cell lines and patient tumors. **(A)** Quantitative RT-PCR of *APOBEC3A* (red) and *APOBEC3B* (blue) mRNA across 34 breast cancer cell lines. N.D., not detected. *n* = 3 biological replicates per line. Values displayed on log₁₀ scale. Individual data points shown. **(B)** *APOBEC3A* versus *APOBEC3B* expression in breast cancer cell lines from DepMap (24Q2 release), log₂(TPM+1). Dashed line, y = x. BT-474 and MDA-MB-453 highlighted. **(C)** Immunoblotting of APOBEC3A and APOBEC3B across a panel of breast cancer cell lines using the 04A04 antibody. Dark and light exposures shown. Vinculin, loading control. **(D–E)** ssDNA cytidine deaminase activity in whole-cell lysates ± RNase A. **(D)** Indicated BT-474 and **(E)** MDA-MB-453 cells. TC-containing oligonucleotide substrate. One-site binding hyperbola fit. Representative of *n* = 3 biological replicates. **(F)** UMAP projections of *APOBEC3C*, *APOBEC3D*, *APOBEC3F*, and *APOBEC3G* expression on the joint BT-474/MDA-MB-453 embedding from Fig. 1B. **(G)** UMAP of the Wu et al. 2021 breast cancer scRNA-seq atlas (GSE176078) colored by *APOBEC3A* expression (left) or cell type annotation (right). **(H)** Percentage of *APOBEC3A*-detected cells per cell type in the Wu et al. dataset. Cell counts indicated. Right panel, mean *APOBEC3A* expression among detected cells per cell type.

**Figure S2.**
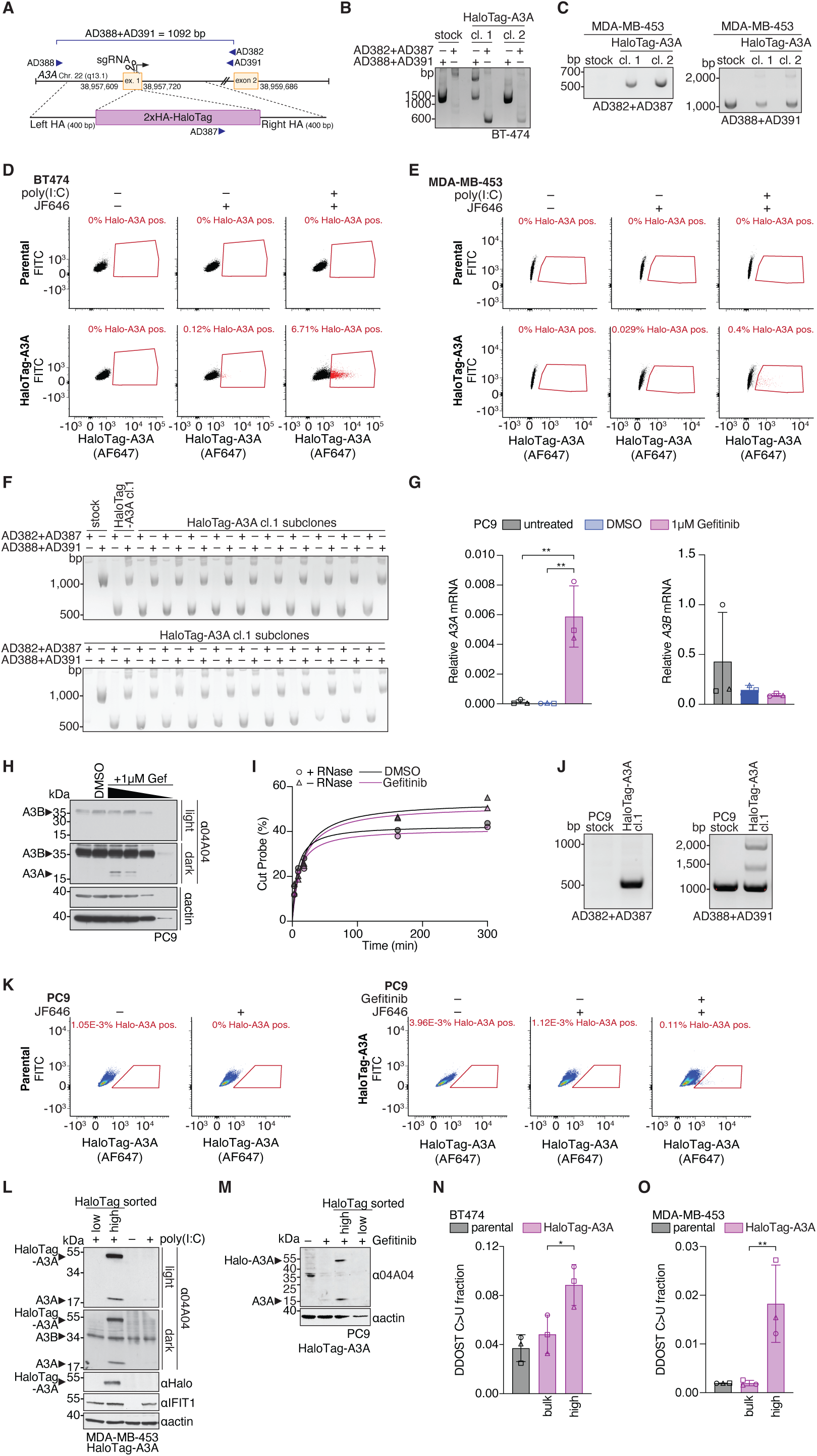
HaloTag-A3A knock-in validation and characterization across three cancer cell lines. **(A)** Targeting schematic for endogenous *APOBEC3A* HaloTag knock-in with PCR primer positions indicated. **(B–C)** PCR validation of HaloTag integration. **(B)** BT-474 (2 clones). **(C)** MDA-MB-453 (2 clones). *n* = 2 experiments. **(D–E)** Representative flow cytometry profiles of HaloTag-A3A labeling ± JF646 ± poly(I:C). **(D)** BT-474. **(E)** MDA-MB-453. Percentage of Halo-A3A-positive cells indicated. **(F)** PCR validation of HaloTag integration across MDA-MB-453 HaloTag-A3A subclones. **(G)** Quantitative RT-PCR of *APOBEC3A* and *APOBEC3B* mRNA in PC-9 cells treated with DMSO or 1 µM gefitinib. *n* = 3 biological replicates. Mean ± s.d. Two-tailed Student’s *t*-test. **(H)** Immunoblotting of APOBEC3A and APOBEC3B in serial dilutions of PC-9 whole-cell lysates ± gefitinib. Actin, loading control. **(I)** ssDNA cytidine deaminase activity in PC-9 whole-cell lysates ± gefitinib ± RNase A. TC-containing substrate. Representative of *n* = 3 biological replicates. **(J)** PCR validation of HaloTag integration in PC-9 HaloTag-A3A clone 1. Two primer pairs shown. **(K)** Representative flow cytometry profiles of PC-9 parental and HaloTag-A3A cells ± JF646 ± gefitinib. Percentage of Halo-A3A-positive cells indicated. **(L)** Immunoblotting of FACS-sorted HaloTag-high and HaloTag-low MDA-MB-453 HaloTag-A3A cells ± poly(I:C). Probed for APOBEC3A (04A04), HaloTag, APOBEC3B (04A04), IFIT1, and actin. Representative of 2 biological replicates. **(M)** Immunoblotting of FACS-sorted HaloTag-high and HaloTag-low PC-9 HaloTag-A3A cells following gefitinib treatment. Probed for APOBEC3A (04A04), HaloTag, and actin. Representative of *n* = 2 biological replicates. **(N–O)** *DDOST* C558 RNA editing (C>U fraction) in FACS-sorted HaloTag-high and unsorted populations. **(N)** BT-474. **(O)** MDA-MB-453. *n* = 3 biological replicates. Mean ± s.d. Student’s *t*-test; \*\**P* < 0.01, \**P* < 0.05. Individual replicate data points overlaid as distinct marker shapes (N and O).

**Figure S3.**
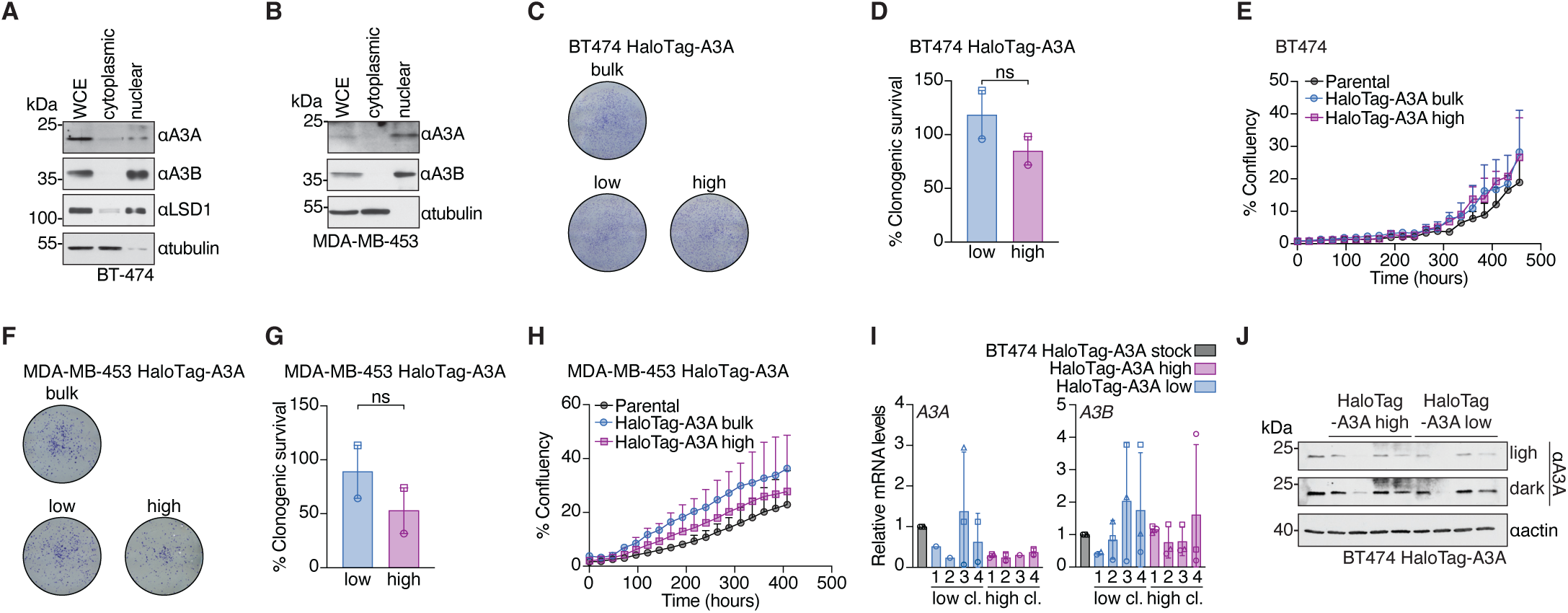
APOBEC3A nuclear localization and transient burst dynamics. (A–B) Subcellular fractionation and immunoblotting. **(A)** BT-474, probed for APOBEC3A (01D05), APOBEC3B, LSD1 (nuclear marker), and tubulin (cytoplasmic marker). **(B)** MDA-MB-453, probed for APOBEC3A, APOBEC3B, and tubulin. Representative of *n* = 2 independent experiments. **(C–D)** Clonogenic survival of FACS-sorted HaloTag-high, HaloTag-low, and bulk BT-474 HaloTag-A3A cells. **(C)** Representative crystal violet staining. **(D)** Quantification. Mean of 2 biological replicates. Student’s *t*-test; ns = not significant. **(E)** Incucyte confluence growth curves of parental BT-474, HaloTag-A3A bulk, and FACS-sorted HaloTag-A3A high cells. Representative of 3 biological replicates. Mean ± s.d. **(F–H)** Same as (C–E) for MDA-MB-453. **(F)** Representative crystal violet staining. **(G)** Clonogenic survival quantification. Mean of 2 biological replicates. Student’s *t*-test; ns = not significant. **(H)** Incucyte confluence growth curves. Representative of 3 biological replicates. Mean ± s.d. **(I)** Quantitative RT-PCR of *APOBEC3A* (left) and *APOBEC3B* (right) mRNA in subclones derived from FACS-sorted HaloTag-high or HaloTag-low BT-474 HaloTag-A3A cells. *n* = 3 independent experiments per subclone. Mean ± s.d. **(J)** Immunoblotting of APOBEC3A (01D05) in subclones derived from FACS-sorted HaloTag-high or HaloTag-low BT-474 HaloTag-A3A cells. Light and dark exposures shown. Actin, loading control. Individual replicate data points overlaid as distinct marker shapes (D, G, and I).

**Figure S4.**
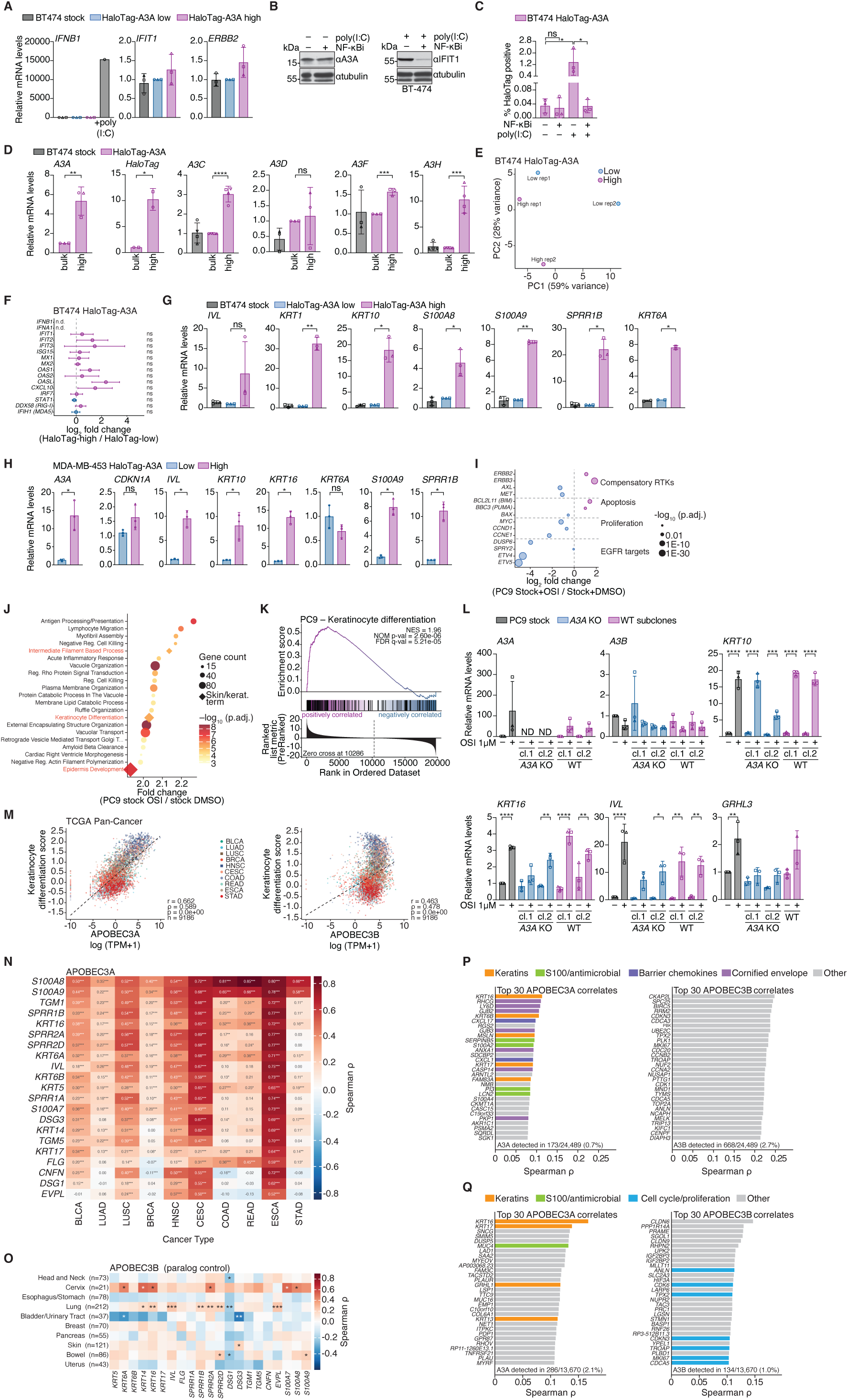
Orthogonal validation of the squamous differentiation state and cross-cancer conservation. **(A)** Quantitative RT-PCR of *IFNB1*, *IFIT1*, and *ERBB2* in BT-474 HaloTag-high and HaloTag-low cells, with poly(I:C)-treated cells as an *IFNB1* positive control. Mean ± s.d. of *n* = 3 independent biological replicates. Statistical analysis by Student’s *t*-test; ns = not significant. **(B)** Immunoblotting with anti-APOBEC3A, anti-IFIT1, and anti-tubulin in BT-474 cells treated with the IKK inhibitor TPCA-1 (1 µM), with or without poly(I:C) stimulation as a positive control for IFIT1 induction. Representative of *n* = 3 independent biological replicates. **(C)** Flow cytometry quantification of HaloTag-A3A high BT-474 cells treated with TPCA-1 (1 µM for 24 hr prior to flow analysis), with or without poly(I:C) stimulation. Mean ± s.d. of *n* = 3 independent biological replicates. Statistical analysis by ANOVA with Tukey’s multiple-comparisons test; \**p* < 0.05, ns = not significant. **(D)** Quantitative RT-PCR of *APOBEC3A*, *HaloTag*, *APOBEC3C*, *APOBEC3D*, *APOBEC3F*, and *APOBEC3H* in BT-474 HaloTag-high and HaloTag-low cells. Mean ± s.d. of *n* = 3 independent biological replicates. Statistical analysis by Student’s *t*-test; \**p* < 0.05, \*\**p* < 0.01, \*\*\**p* < 0.001, \*\*\*\**p* < 0.0001, ns = not significant. **(E)** Principal component analysis of BT-474 HaloTag-high and HaloTag-low RNA-seq samples. **(F)** Forest plot of log₂ fold change and DESeq2 log₂ fold change standard error (lfcSE) for selected IFN/ISG transcripts in the BT-474 HaloTag-high vs HaloTag-low RNA-seq data. Error bars represent lfcSE. No genes shown crossed *p*adj < 0.05. n.d., not detected. **(G–H)** Quantitative RT-PCR validation of squamous markers in **(G)** BT-474 stock, HaloTag-high, and HaloTag-low cells, and **(H)** poly(I:C)-stimulated MDA-MB-453 HaloTag-high and HaloTag-low cells. Mean ± s.d. of *n* = 3 independent biological replicates. Statistical analysis by two-tailed Student’s *t*-test; \**p* < 0.05, \*\**p* < 0.01, \*\*\**p* < 0.001, ns = not significant. **(I)** Dot plot of log₂ fold change (WT + OSI vs WT + UT) for compensatory receptor tyrosine kinases, apoptosis markers, proliferation drivers, and EGFR pathway targets. Dot size, −log₁₀(*p*adj) on a log scale; red, upregulated; blue, downregulated. DESeq2 Wald test, Benjamini-Hochberg FDR. **(J)** Gene ontology biological process bubble plot of pathways enriched in the osimertinib response of WT PC-9 cells (fgsea, ranked by DESeq2 Wald statistic, 10,000 permutations, Benjamini-Hochberg FDR). Skin and keratinization terms highlighted with red diamonds. Bubble size, leading-edge gene count; color, −log₁₀(*p*adj). **(K)** Gene set enrichment analysis of GO_KERATINOCYTE_DIFFERENTIATION (GO:0030216, 112 genes after size filtering) in the osimertinib response of *APOBEC3A* knockout PC-9 cells. Ranked metric, DESeq2 Wald statistic. NES = 1.96, NOM *p* = 2.6 × 10⁻⁶, FDR *q* = 5.2 × 10⁻⁵ (fgsea, 10,000 permutations). **(L)** Quantitative RT-PCR of *APOBEC3A*, *APOBEC3B*, *KRT10*, *KRT16*, *IVL*, and *GRHL3* in PC-9 stock, two independent *APOBEC3A* knockout clones, and two WT subclones, treated with DMSO or 1 µM osimertinib. Mean ± s.d. of *n* = 3 independent biological replicates. Statistical analysis by two-way ANOVA with Tukey’s multiple-comparisons test; \**p* < 0.05, \*\**p* < 0.01, \*\*\**p* < 0.001, \*\*\*\**p* < 0.0001, ns = not significant. n.d., not detected. **(M)** Pan-cancer TCGA scatter of *APOBEC3A* (left) and *APOBEC3B* (right) expression versus a 21-gene keratinocyte differentiation score (mean z-score of *KRT5*, *KRT6A*, *KRT6B*, *KRT14*, *KRT16*, *KRT17*, *IVL*, *FLG*, *SPRR1A*, *SPRR1B*, *SPRR2A*, *SPRR2D*, *DSG1*, *DSG3*, *TGM1*, *TGM5*, *CNFN*, *EVPL*, *S100A7*, *S100A8*, *S100A9*). *n* = 9,186 primary tumours across 32 TCGA cancer types; 10 APOBEC-relevant cancer types are highlighted. Pearson and Spearman correlations are reported on each panel. The comparison of *APOBEC3A* versus *APOBEC3B* correlations used a Meng-Rosenthal-Rubin test for dependent correlations (*p* = 1.1 × 10⁻⁴⁰). **(N)** Heatmap of per-gene, per-cancer-type Spearman correlations between *APOBEC3A* expression and individual keratinocyte signature genes across TCGA cohorts. Significance thresholds based on Benjamini-Hochberg FDR across all gene × cancer-type pairs: *FDR < 0.05, **FDR < 0.01, ***FDR < 0.001. Squamous-enriched cancer types (LUSC, HNSC) marked with filled squares. **(O)** Heatmap of per-gene, per-lineage Spearman correlations between *APOBEC3B* expression and individual keratinocyte signature genes across DepMap cancer cell lines. Significance: \**p* < 0.05, \*\**p* < 0.01, \*\*\**p* < 0.001. DepMap 24Q2. Shown as paralog control for Fig. 2K. Meng-Rosenthal-Rubin test for *APOBEC3A* vs *APOBEC3B* correlations, *p* < 10⁻¹⁵. **(P)** Top 30 transcriptional correlates of *APOBEC3A* (left) and *APOBEC3B* (right) in the Wu et al. 2021 breast cancer scRNA-seq atlas. *n* = 24,489 cancer epithelial cells from 26 patients. Bars coloured by functional category. Spearman correlation with Benjamini-Hochberg FDR across 10,634 tested genes; only hits with ρ > 0.05 and FDR < 0.05 shown. **(Q)** Top 30 transcriptional correlates of *APOBEC3A* (left) and *APOBEC3B* (right) in the Kim et al. 2020 lung adenocarcinoma scRNA-seq atlas. *n* = 13,670 primary LUAD malignant cells (metastatic and non-malignant samples excluded). Bars coloured by functional category. Spearman correlation with Benjamini-Hochberg FDR; only hits with ρ > 0.05 and FDR < 0.05 shown. Individual replicate data points overlaid as distinct marker shapes (A, C, D, G, H, and L).

**Figure S5.**
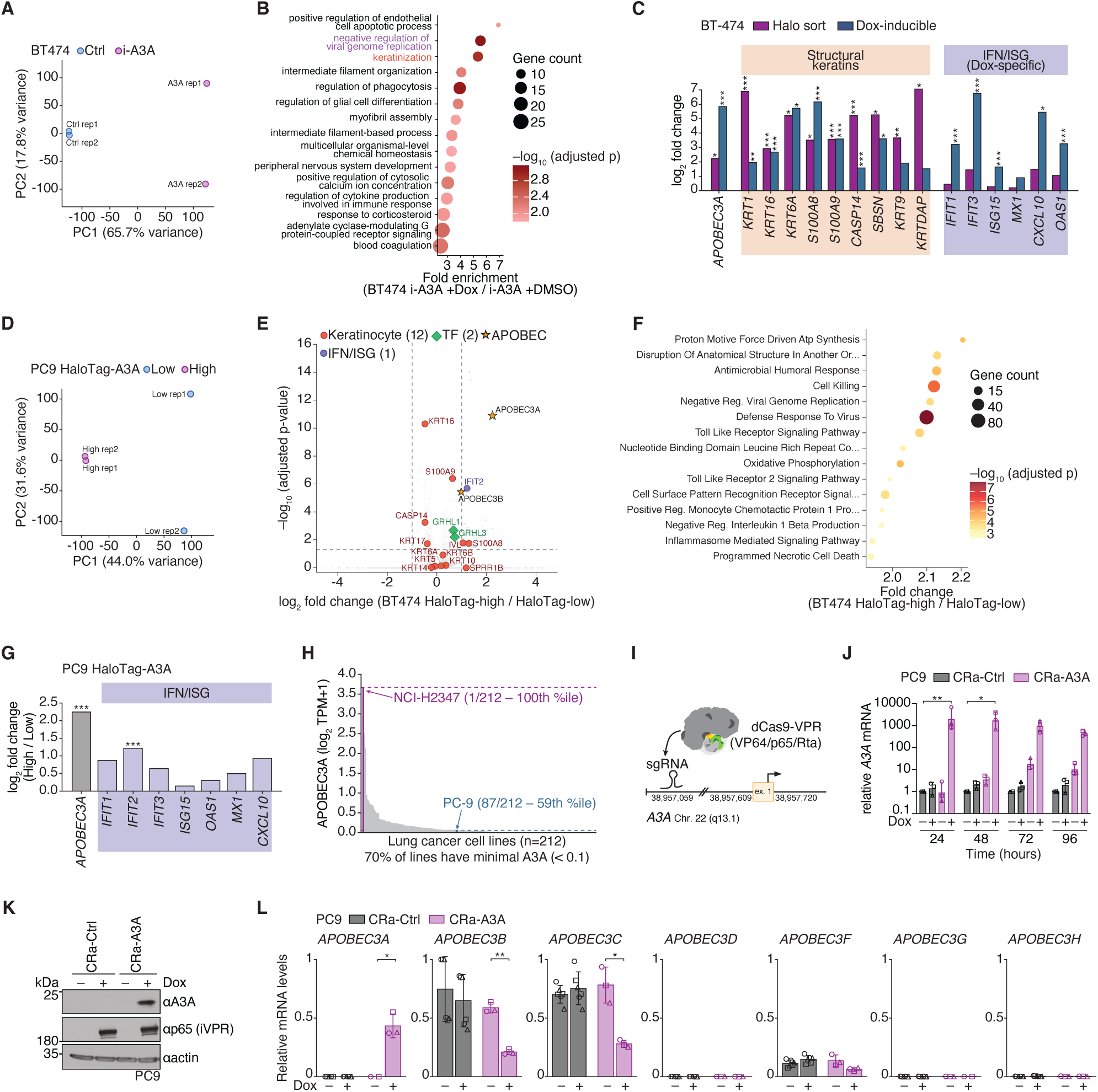
Extended characterization of APOBEC3A-driven transcriptional responses in BT-474 and PC-9 cells. **(A)** Principal component analysis of BT-474 cells with dox-inducible APOBEC3A, induced (A3A) or uninduced (Ctrl). PC1, 65.7% variance; PC2, 17.8% variance. *n* = 2 biological replicates per condition. PCA on DESeq2 peak-normalized counts after filtering genes with mean > 1, log₂ transformation, and per-gene standardization. **(B)** Gene ontology (biological process) over-representation analysis of upregulated DEGs (*p*adj < 0.05 and log₂FC > 1) in dox-induced BT-474 cells. clusterProfiler::enrichGO with Benjamini-Hochberg adjustment and simplification (cutoff = 0.6). Top 15 terms by fold enrichment. Dot size, gene count. Color, −log₁₀ adjusted *p*-value. **(C)** Log₂ fold change of ten concordantly regulated keratinocyte and interferon-stimulated genes in BT-474 Halo-sort (orange) and Dox-OE (red) datasets. DESeq2 Wald test. **p*adj < 0.05, \*\**p*adj < 0.01, \*\*\**p*adj < 0.001. **(D)** Principal component analysis of PC-9 Halo-A3A-high and Halo-A3A-low cells. PC1, 44.0% variance; PC2, 31.6% variance. *n* = 2 biological replicates per condition. Normalization and processing as in (A). **(E)** Volcano plot of differential expression in PC-9 Halo-A3A-high vs Halo-A3A-low cells. *n* = 2 biological replicates per condition. DESeq2 Wald test. Significance thresholds, *p*adj < 0.05 and |log₂FC| > 1. Color coding in the panel. Labeled genes have baseMean ≥ 5. **(F)** Gene set enrichment analysis of GO biological process terms in PC-9 Halo-A3A-high vs Halo-A3A-low cells. fgsea preranked on DESeq2 Wald statistic, minimum/maximum set size 10/500, 10,000 permutations. Top 15 terms by NES after Jaccard leading-edge redundancy filter. Bubble size, leading-edge gene count. Color, −log₁₀ adjusted *p*-value. **(G)** Log₂ fold change of canonical interferon-stimulated genes in PC-9 Halo-A3A-high vs Halo-A3A-low cells. *APOBEC3A* (orange) is shown as a positive control for the sort. *n* = 2 biological replicates per condition. DESeq2 Wald test. \*\*\**p*adj < 0.001, ns = not significant. **(H)** *APOBEC3A* expression across 212 lung cancer cell lines (DepMap 24Q2 release), ranked by log₂(TPM+1). NCI-H2347 (red) and PC-9 (blue) are indicated. **(I)** Schematic of CRISPR activation at the endogenous *APOBEC3A* locus. dCas9-VPR is recruited by a single-guide RNA targeting the *APOBEC3A* promoter. **(J)** Quantitative RT-PCR of *APOBEC3A* mRNA in PC-9 dCas9-VPR cells expressing control or *APOBEC3A*-targeting sgRNA across a 96 h doxycycline time course. *n* = 3 biological replicates. One-way ANOVA with Tukey’s multiple comparisons. Error bars, mean ± s.d. \**p* < 0.05, \*\**p* < 0.01. **(K)** Immunoblot of APOBEC3A, iVPR (p65), and actin in PC-9 dCas9-VPR cells expressing control or *APOBEC3A*-targeting sgRNA, with or without doxycycline. Representative of *n* = 3 biological replicates. **(L)** Quantitative RT-PCR of *APOBEC3B*, *APOBEC3C*, *APOBEC3D*, *APOBEC3F*, *APOBEC3G*, and *APOBEC3H* in PC-9 dCas9-VPR cells expressing two control sgRNAs (CTRLsg1, CTRLsg3) or two *APOBEC3A*-targeting sgRNAs (sg28, sg28/30), with or without doxycycline. *n* = 3 biological replicates. Error bars, mean ± s.d. Individual replicate data points overlaid as distinct marker shapes (J and L).

**Figure S6.**
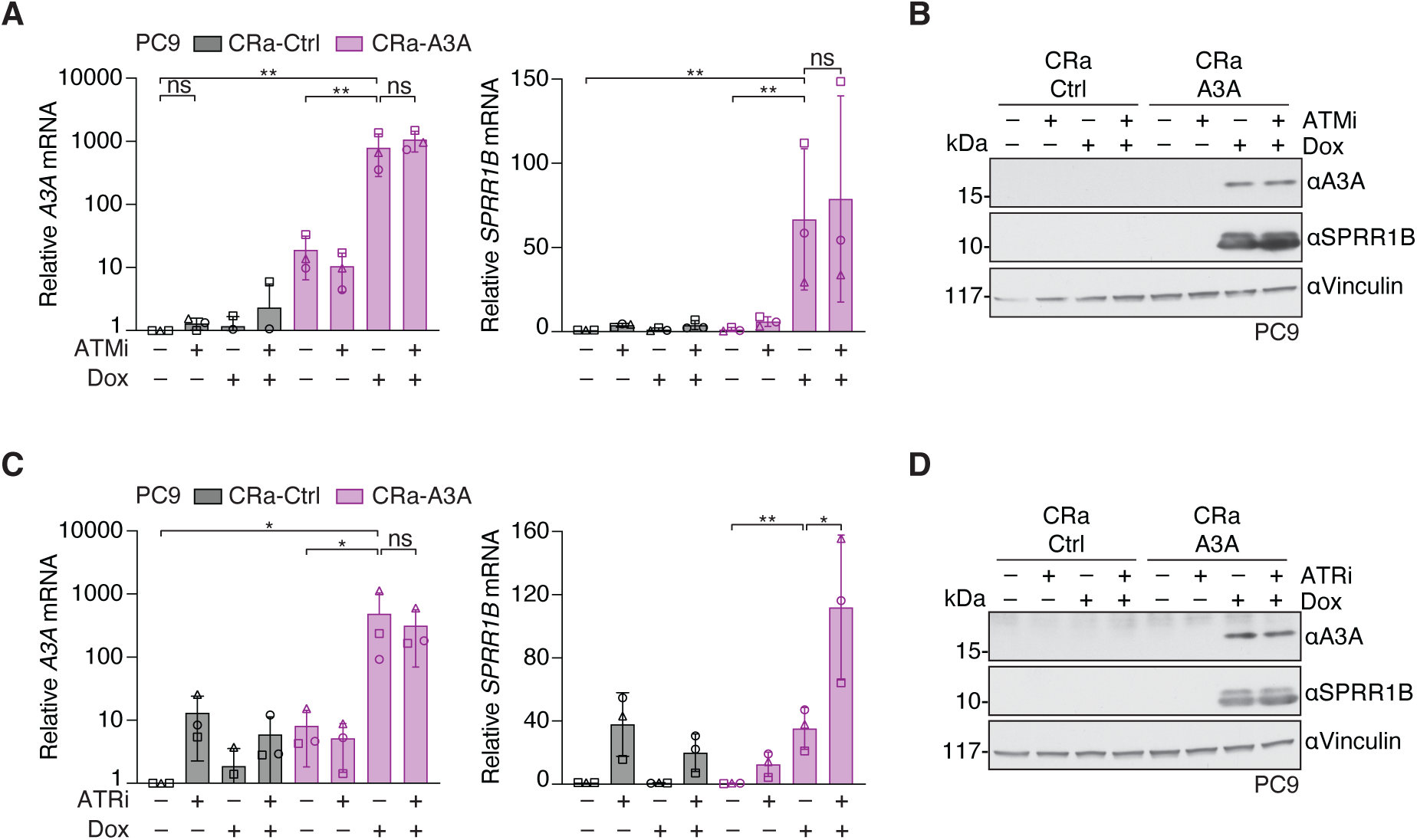
ATM and ATR are dispensable for APOBEC3A-driven gene induction. **(A)** Quantitative RT-PCR of *APOBEC3A* and *SPRR1B* in PC-9 dCas9-VPR cells expressing CRa-Ctrl or CRa-A3A, treated with doxycycline (1 μg/mL) and/or ATM inhibitor (10 μM). *n* = 3 independent biological replicates. One-way ANOVA with Tukey’s multiple-comparisons test. *APOBEC3A* values displayed on log₁₀ scale. Error bars, mean ± s.d. *p*-values as indicated; ns, not significant. **(B)** Immunoblot of APOBEC3A, SPRR1B, and vinculin in PC-9 dCas9-VPR cells expressing CRa-Ctrl or CRa-A3A, treated with doxycycline (1 μg/mL) and/or ATM inhibitor (10 μM). Representative of *n* = 1 biological experiment. Antibodies listed in the methods section. **(C)** Quantitative RT-PCR of *APOBEC3A* and *SPRR1B* in PC-9 dCas9-VPR cells expressing CRa-Ctrl or CRa-A3A, treated with doxycycline (1 μg/ mL) and/or ATR inhibitor (5 μM). *n* = 3 independent biological replicates. One-way ANOVA with Tukey’s multiple-comparisons test. *APOBEC3A* values displayed on log₁₀ scale. Error bars, mean ± s.d. *p*-values as indicated; ns, not significant. **(D)** Immunoblot of APOBEC3A, SPRR1B, and vinculin in PC-9 dCas9-VPR cells expressing CRa-Ctrl or CRa-A3A, treated with doxycycline (1 μg/mL) and/or ATR inhibitor (5 μM). Representative of *n* = 1 biological experiment. Antibodies listed in the methods section. Individual replicate data points overlaid as distinct marker shapes (A and C).

**Figure S7.**
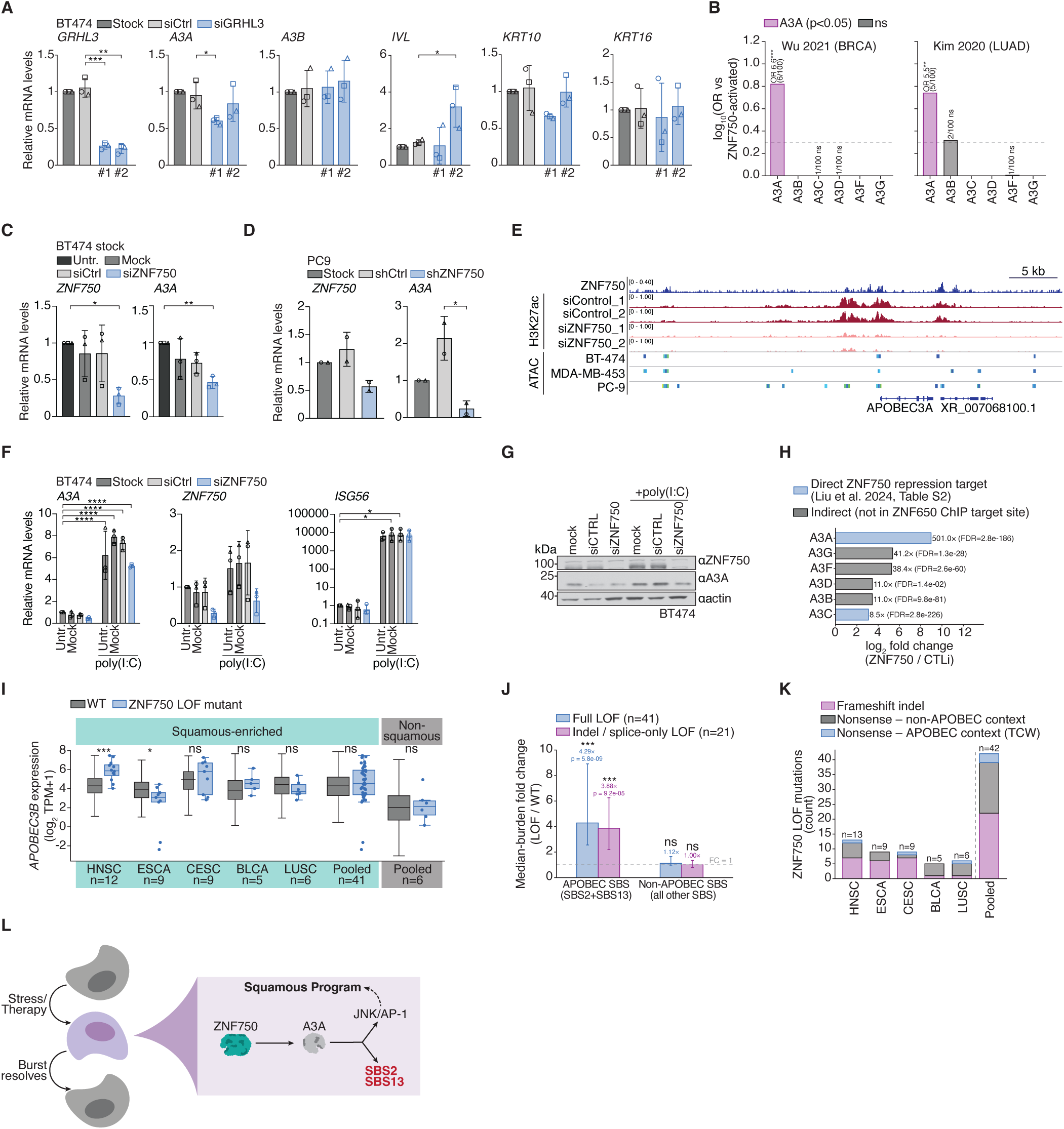
ZNF750 regulation of APOBEC3A across cancer lineages and squamous tumors. **(A)** Quantitative RT-PCR of *GRHL3*, *APOBEC3A*, *APOBEC3B*, *IVL*, *KRT10*, and *KRT16* in BT-474 Stock, siCtrl, and siGRHL3 (#1 and #2) cells. *n* = 3 independent biological replicates. Mean ± s.d. Unpaired two-tailed *t*-test. \**p* < 0.05, \*\**p* < 0.01, \*\*\**p* < 0.001. Individual replicate data points overlaid as distinct marker shapes. **(B)** Enrichment of ZNF750-activated targets among the top 100 transcriptional correlates of each APOBEC3 paralog in patient tumor scRNA-seq. Left, Wu 2021 breast cancer (GSE176078). Right, Kim 2020 lung adenocarcinoma (GSE131907). Odds ratio by Fisher’s exact test. Dashed line, OR = 2. \*\**p* < 0.01, \*\*\**p* < 0.001, ns = not significant. **(C)** Quantitative RT-PCR of *ZNF750* and *APOBEC3A* in BT-474 Stock, siCtrl, and siZNF750 cells (untransfected and mock controls). *n* = 3 independent biological replicates. Mean ± s.d. Unpaired two-tailed *t*-test. \**p* < 0.05, \*\**p* < 0.01. Individual replicate data points overlaid as distinct marker shapes. **(D)** Quantitative RT-PCR of *ZNF750* and *APOBEC3A* in PC-9 Stock, shCtrl, and shZNF750 cells. *n* = 3 independent biological replicates. Mean ± s.d. Unpaired two-tailed *t*-test. \**p* < 0.05. **(E)** Genome browser view of the *APOBEC3A* locus. Top, ZNF750 ChIP-seq in differentiated keratinocytes (Boxer 2014, GSE57702). Middle, H3K27ac ChIP-seq in siControl and siZNF750 keratinocytes (GSE84657). Bottom, ATAC-seq peaks in BT-474, MDA-MB-453, and PC-9 cells. Scale bar, 5 kb. **(F)** Quantitative RT-PCR of *APOBEC3A*, *ZNF750*, and *ISG56* in BT-474 Stock, siCtrl, and siZNF750 cells ± poly(I:C). *n* = 3 independent biological replicates. Mean ± s.d. Unpaired two-tailed *t*-test. \**p* < 0.05, \*\*\*\**p* < 0.0001. Individual replicate data points overlaid as distinct marker shapes. **(G)** Immunoblot of ZNF750, APOBEC3A, and actin in BT-474 mock, siCTRL, and siZNF750 cells ± poly(I:C). Representative of *n* = 2 independent biological replicates. Antibodies listed in the methods. **(H)** APOBEC3 paralog expression upon *ZNF750* depletion in differentiated keratinocytes (reanalyzed from Liu et al. 2024, GSE248486). Blue, direct ZNF750 ChIP target. Grey, indirect. Linear fold change and FDR indicated. **(I)** *APOBEC3B* mRNA expression (log₂(TPM+1)) in *ZNF750* LOF versus WT tumors across five TCGA squamous-enriched cancer types and a pooled non-squamous cohort. Box plots show median, interquartile range, and 1.5× IQR whiskers. BH-adjusted Mann-Whitney U test. \**p* < 0.05, \*\*\**p* < 0.001, ns = not significant. *n* per group indicated. **(J)** Median-burden fold change (LOF/WT) for APOBEC SBS (SBS2+SBS13) and non-APOBEC SBS in *ZNF750* LOF tumors across TCGA squamous cancers. Full LOF (*n* = 40) and indel/ splice-only LOF (*n* = 21) shown separately. Error bars, 95% bootstrap CI. Dashed line, FC = 1. \*\*\**p* < 0.001, ns = not significant. **(K)** Mutational spectrum of *ZNF750* LOF events across TCGA squamous cancer types. Frameshift indels (pink), nonsense mutations in non-APOBEC context (grey), and nonsense mutations in APOBEC trinucleotide context (TCW, blue). *n* per cancer type indicated. **(L)** Model of squamous-state excursions and APOBEC3A activation in cancer. Stress or therapy induces rare cancer cells to enter a transient squamous differentiation state. ZNF750 supports APOBEC3A induction within this state. APOBEC3A catalytic activity reinforces selected components of the squamous program through uracil excision and JNK/AP-1 signaling, linking transient lineage-state plasticity to SBS2/SBS13 mutational processes. The APOBEC3A-high excursion resolves over days.

## SUPPLEMENTARY TABLE LEGENDS

**Table S1.** Differential gene expression in BT-474 HaloTag-A3A-high versus HaloTag-A3A-low cells. DESeq2 results for 22,147 genes. Columns report baseMean, log_2_ fold change, log_2_ fold change standard error, Wald statistic, nominal p-value, and Benjamini-Hochberg adjusted p-value. n = 2 biological replicates per condition.

**Table S2.** Gene ontology biological process over-representation analysis of BT-474 Halo-sort upregulated DEGs. clusterProfiler enrichGO output for 17 upregulated genes (padj < 0.05, log_2_FC > 0) tested against 12,888 Entrez-mapped background genes. Redundant terms collapsed using simplify (cutoff = 0.6). 76 enriched terms reported.

**Table S3.** Gene set enrichment analysis results. Sheets 1-7 report fgsea results (v1.36.2) for seven PC-9 and BT-474 contrasts using MSigDB 2026.1.Hs C5 GO biological process gene sets (minSize = 10, maxSize = 500, 10,000 permutations). Sheets 8-9 report Broad GSEA v4.3.2 GseaPreranked results for BT-474 Halo-sort using MSigDB v6.0 C5 GO gene sets (classic enrichment statistic, 1,000 permutations). Columns report pathway name, normalized enrichment score, nominal p-value, adjusted p-value, gene set size, and leading-edge genes.

**Table S4.** Differential gene expression in PC-9 cells across three contrasts. Sheet 1, wild-type osimertinib-treated versus untreated (19,393 genes). Sheet 2, *APOBEC3A* knockout osimertinib-treated versus untreated (19,721 genes). Sheet 3, HaloTag-A3A-high versus HaloTag-A3A-low under osimertinib (21,281 genes). DESeq2. n = 2 biological replicates per condition. Column structure as in Table S1.

**Table S5.** Differential gene expression in BT-474 doxycycline-induced *APOBEC3A* versus uninduced controls. DESeq2 results for 23,121 genes. Column structure as in Table S1.

**Table S6.** Guide RNA and shRNA sequences used in this study.

**Table S7.** Antibodies used in this study.

**Table S8.** Primer sequences used in this study.

